# Multiple separate cases of pseudogenized meiosis genes Msh4 and Msh5 in Eurotiomycete fungi: associations with Zip3 sequence evolution and homothallism, but not Pch2 losses

**DOI:** 10.1101/750497

**Authors:** Elizabeth Savelkoul, Cynthia Toll, Nathan Benassi, John M. Logsdon

## Abstract

The overall process of meiosis is conserved in many species, including some lineages that have lost various ancestrally present meiosis genes. The extent to which individual meiosis gene losses are independent from or dependent on one another is largely unknown. Various Eurotiomycete fungi were investigated as a case system of recent meiosis gene losses after BLAST and synteny comparisons found Msh4, Msh5, Pch2, and Zip3 to be either pseudogenized or undetected in *Aspergillus nidulans* yet intact in congeners such as *A. fumigatus*. Flanking gene-targeted degenerate PCR primers applied to 9 additional *Aspergillus* species found (i) Msh4, Msh5, and Zip3 pseudogenized in *A. rugulosus* (sister taxon to *A. nidulans*) but intact in all other amplified sequences; and (ii) Pch2 not present at the syntenic locus in most of the 9 species. Topology tests suggested two independent Pch2 losses in genus *Aspergillus*, neither directly coinciding with pseudogenization of the other three genes. The *A. nidulans-A. conjunctus* clade Pch2 loss was not associated with significant K_a_/K_s_ changes for Msh4, Msh5, or Zip3; this suggests against prior Pch2 loss directly altering sequence evolution constraints on these three genes. By contrast, Zip3 K_a_/K_s_ tended to be elevated in several other Eurotiomycete fungi with independently pseudogenized Msh4 and Msh5 (*Talaromyces stipitatus*, *Eurotium herbariorum*). The coinciding K_a_/K_s_ elevation and/or clear pseudogenization of Zip3 in taxa with pseudogenized Msh4 and Msh5 is consistent with some degree of molecular coevolution. Possible molecular, environmental, and life history variables (*e.g.*, homothallism) that may be associated with these numerous independent meiosis gene losses (Msh4: 3, Msh5: 3, Zip3: ≥ 1, Pch2: 4) are discussed.

## Introduction

Meiosis, a form of cell division that includes both ploidy reduction and elevated recombination rates between homologous chromosomes, is essential for successful sexual reproduction in many species. All major eukaryotic supergroups include at least some sexual taxa, many of which also have conserved orthologs of genes encoding proteins with established functions in meiosis (“meiosis genes”) or exclusively affecting meiosis (“meiosis-specific genes”); this combination of conserved presence and conserved function across many lineages is consistent with the ancestral state of eukaryotes having a core set of meiosis genes both present and necessary for successful meiosis (Ramesh *et al*., 2005; Malik *et al*., 2008; Schurko and Logsdon, 2008). Meiosis-specific genes are often identified by knockout mutant phenotypes that confer a sterility or reduced fertility phenotype in the examined taxon with no other somatic or vegetative growth effects. However, several established sexual model taxa (*e.g.*, *Schizosaccharomyces pombe, Caenorhabditis elegans, Drosophila melanogaster*) are capable of successful meiosis despite lacking detectable orthologs of a subset of meiosis genes essential for viable gamete or spore production in other taxa (Malik *et al*., 2008). This presents a mechanistic puzzle: how have some lineages successfully transitioned from an ancestral state where loss of function in a gene compromises meiosis to a derived state where successful meiosis is maintained despite loss of the gene?

One obstacle to identifying the most influential factors for these transitions is that most previously reported meiosis gene losses have been relatively ancient (*i.e.*, no detectable orthologs and/or no close relatives known to retain the genes) and obtained from investigations that focus on breadth of taxon sampling (*i.e.*, a few exemplar taxa from multiple diverse eukaryotic lineages; *e.g.*, Ramesh *et al*., 2005; Malik *et al*., 2008). By contrast, identification of relatively more recent meiosis gene losses (*e.g.*, with still-detectable pseudogenes) within a group of closely related sexual taxa that include some species retaining the genes would be more amenable to comparative biology studies.

Fungi have been the subjects of an increasing number of taxonomically broad (Malik *et al*., 2008; Halary *et al*., 2011) or focused (Galagan *et al*., 2005; Woo *et al*., 2006; Butler *et al*., 2009; Desjardins *et al*., 2011; Martinez *et al*., 2012) surveys of meiosis gene distribution. Many of these efforts have been in the context of assessing species’ potential for cryptic sexual reproduction, reasoning that genes encoding proteins that function exclusively in meiosis would be expected to pseudogenize and degrade in obligately asexual taxa yet retained intact if sexual reproduction still occurs (Schurko and Logsdon, 2008). Pathogenic fungi in class Eurotiomycetes of phylum Ascomycota (*e.g.*, *Coccidioides* spp.*, Trichophyton* spp., *Microsporum* spp. (Cox and Magee, 2004; Martinez *et al*., 2012)) have been a particular subject of interest for two reasons: first, the conditions under which these species can be induced to undergo sexual reproduction were unknown for many years (Horn *et al*., 2009a; Horn *et al*., 2009b; O’Gorman *et al*., 2009) or remain unknown (*Paracoccidioides brasiliensis* (Matute et al., 2006); *Aspergillus niger, Trichophyton* spp., *Coccidioides* spp. (Broad, 2012)); second, higher-virulence genotypes could arise through the genetic variation produced during sexual reproduction *via* meiosis (McDonald and Linde, 2002; Li *et al*., 2012). Previously examined Eurotiomycetes, primarily from order Onygenales (*e.g., Coccidioides* and *Trichophyton* spp.) but also some *Aspergillus* species from order Eurotiales (Wang *et al*., 2009), have either no reported cases of meiosis-specific gene losses (Malik *et al*., 2008; Martinez *et al*., 2012) or sporadic instances of possible meiosis-specific gene losses (Woo *et al*., 2006; Desjardins *et al*., 2011). However, our present analyses of numerous additional Eurotiomycete genomes by bioinformatics and PCR have found that the previously reported general conservation of meiosis-specific genes in these fungi has several striking exceptions in order Eurotiales—including confirmation of recent pseudogenization of several meiosis genes in the model sexual Eurotiomycete *Aspergillus nidulans* (Todd *et al*., 2007).

While conducting a broader TBLASTN-based bioinformatics survey of meiosis genes across diverse fungal lineages with sequenced genomes (Savelkoul, 2013; relevant subsections included in this work), *Aspergillus nidulans* was found to have two major differences relative to other available sequenced *Aspergillus*: (i) an ortholog of the pachytene checkpoint gene Pch2 (San-Segundo and Roeder, 1999) was undetected in *A. nidulans* and (ii) three “ZMM” group crossover resolution genes—Msh4, Msh5, and Zip3 (Lynn *et al*., 2007)—were pseudogenized and in various states of degradation in *A. nidulans*. Two major questions became apparent. First, have other Eurotiomycetes also lost any or all of these four genes? Second, are the losses of these four genes functionally related to each other? To investigate these questions, we characterized Msh4, Msh5, Zip3, and Pch2 phylogenetic distributions and evolutionary rates (K_a_/K_s_) using publicly available Eurotiomycete genome sequences and degenerate PCR on additional *Aspergillus* species lacking sequenced genomes. Our results are consistent with multiple relatively recent independent losses of Msh4, Msh5, and Pch2, as well as at least one loss of Zip3, in the examined taxa; these indicate a previously undescribed tendency for convergent alterations to meiotic crossover formation pathways among Eurotiales fungi.

## Results

### Initial Bioinformatic Inventory and Synteny Assessment

TBLASTN searches of several initially examined Eurotiomycete genome sequences (*Aspergillus nidulans, Aspergillus terreus, Aspergillus flavus, Aspergillus niger, Aspergillus clavatus, Aspergillus fumigatus, Coccidioides immitis, Uncinocarpus reesii, Histoplasma capsulatum, Paracoccidioides brasiliensis, Microsporum gypseum, Microsporum canis, Trichophyton rubrum, Trichophyton tonsurans*; see supplemental file S1 for genome providers and strain information) successfully identified putative orthologs of Msh4, Msh5, and Pch2 in all species except *A. nidulans* (Figure 1, see supplemental file S1 for accession numbers). The putative orthologs identified at this initial stage appeared to be intact (*i.e.*, no in-frame stop codons, loss of consensus splice site sequences in introns with highly conserved positions, frameshifts, or large deletions.) No *A. nidulans* Msh4, Msh5, or Pch2 orthologs were identified by TBLASTN searches despite the search methods being sufficient to (i) identify these orthologs in congeners and (ii) return paralogous hits in the *A. nidulans* genome (Msh1, Msh2, Msh3, Msh6; other AAA ATPases; data not shown). Synteny around Msh4, Msh5, and Pch2 was generally well-conserved in these Eurotiomycetes; the consensus was Msh4 flanked by the WD repeat-containing ribosome biogenesis protein Rsa4 (“Rsa4”) and a conserved C2H2 finger domain protein (“C2H2”), Msh5 flanked by heat shock protein Hsp78 (“Hsp78”) and a Sas10/Utp3 family protein (“Sas10”), and Pch2 flanked by the secretion pathway F-box protein Pof6/Sls2/Rcy1 (“Rcy1”) and cytochrome C1/Cyt1 (“Cyt1”) (Figure 2, see supplemental file S1 for accession numbers). These flanking gene pairs (Rsa4 and C2H2, Hsp78 and Sas10, Rcy1 and Cyt1) were then successfully identified in the *A. nidulans* genome, where they were retained in the Eurotiomycete consensus orientations (Figure 2). The *A. nidulans* intergenic regions (Rsa4-C2H2: 696 bp, Hsp78-Sas10: 1067 bp, Rcy1-Cyt1: 1063 bp) were notably shorter than exemplar full-length *A. clavatus* orthologs of Msh4 (2958 bp), Msh5 (3702 bp), and Pch2 (1778 bp). Nevertheless, highly degraded pseudogenes of Msh4 and Msh5 were identified in the *A. nidulans* syntenic locations (Figure 2, Table 1) that could be aligned with intact orthologs (Figure 3A, Figure 3B) and showed the phylogenetic placement expected for *A. nidulans* relative to other *Aspergillus* species (Figure 4, Figure 5). Neither manual inspection nor BLAST2SEQ comparisons to intact *Aspergillus* Pch2 orthologs yielded recognizable, alignable residual Pch2 sequence in the *A. nidulans* Rcy1-Cyt1 intergenic region (data not shown).

**Figure 1:**
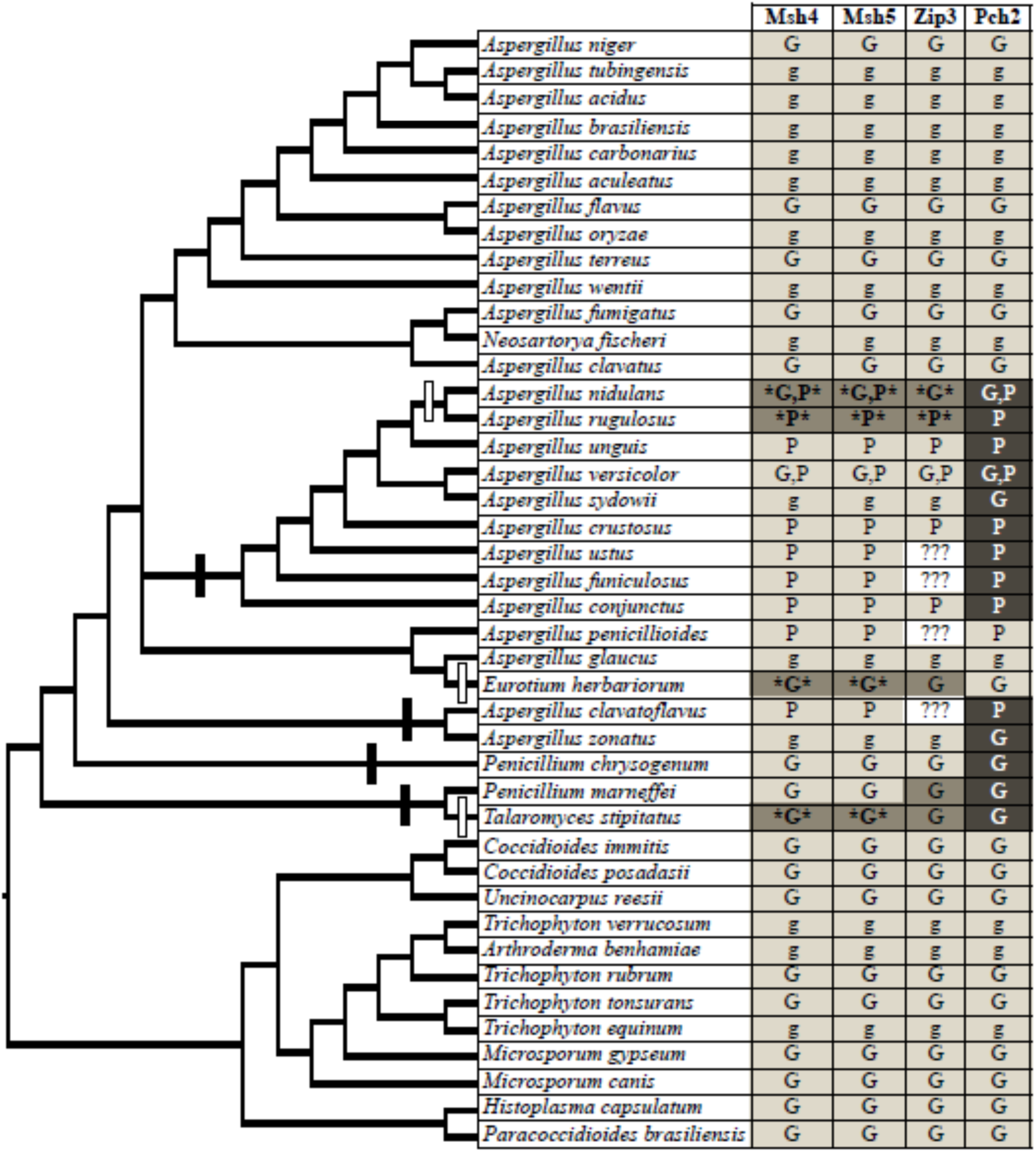
Summary of Msh4, Msh5, Pch2, and Zip3 Distribution in Examined Eurotiomycetes. Sequence sources are indicated by “G” (BLAST searches of publicly available genome sequences) or “P” (PCR in this work). Grid square colors and text formatting indicate ortholog status: present and apparently intact (light gray with lower case = manual annotation only, light gray with upper case = manual annotation and phylogenetic validation); possible pseudogenes (medium gray, plain font); pseudogenes (medium gray, bold font with * *); ortholog not detected by genome TBLASTN searches or at syntenic genome region amplified by PCR (dark gray, white font); data not available due to PCR amplification failure (white, “???”). Putative independent gene loss events are noted on the cladogram with black rectangles (Pch2) and white rectangles (Msh4, Msh5). The phylogeny is derived from Hubka *et al*. (2013), Martinez *et al*. (2012), Peterson (2008), van den Berg *et al*. (2008), and Wang *et al*. (2009) with additional consultation from the JGI *Aspergillus acidus* genome home page (JGI, 2014) and NCBI Taxonomy database (NCBI, 2014).

**Figure 2:**
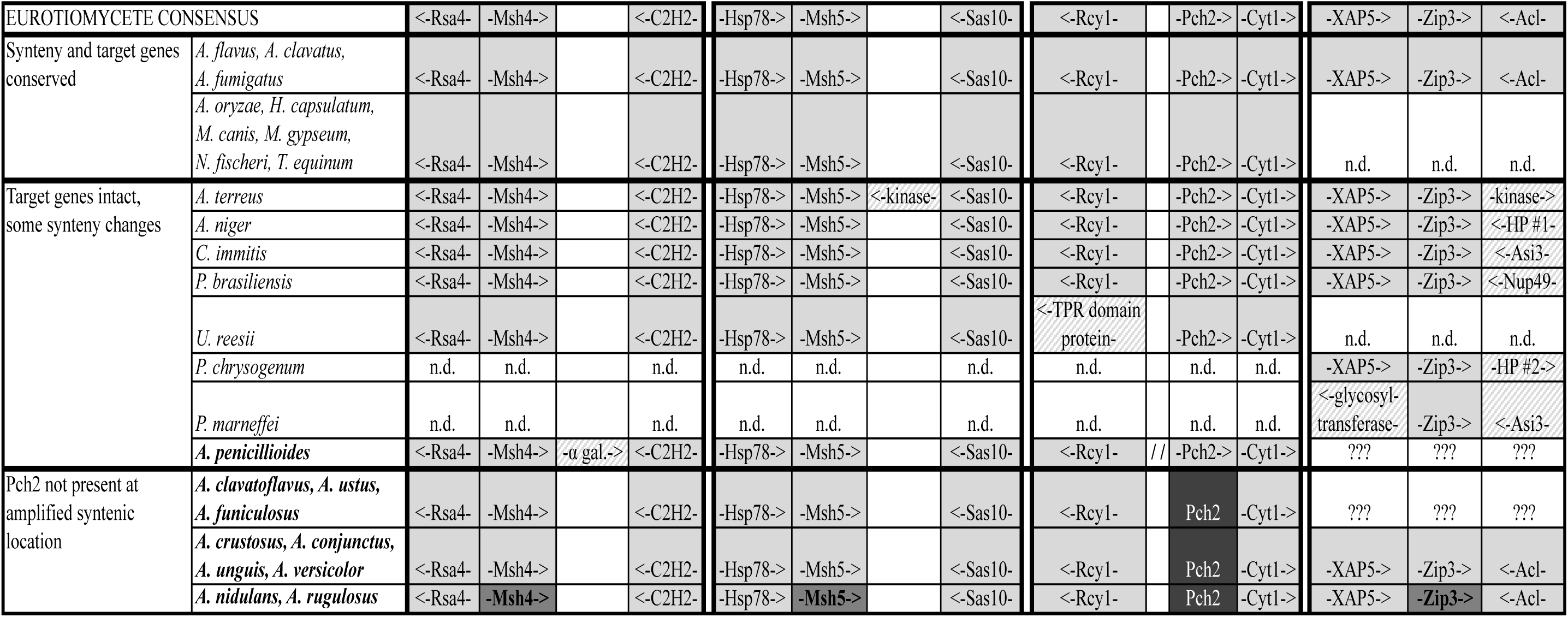
Eurotiomycete Synteny Surrounding Msh4, Msh5, Pch2, and Zip3. Species used to determine consensus synteny surrounding Msh4, Msh5, Pch2, and Zip3 in Eurotiomycetes are listed in plain text; species subjected to degenerate PCR are in bold font. Species with identical gene distribution patterns are collapsed into a single row. Solid light gray boxes are orthologs matching the consensus synteny that are present and apparently intact. Hatched light gray and white boxes are apparently intact genes that do not follow consensus synteny. Solid medium gray boxes with bold font are pseudogenes. Dark gray boxes with white font indicate no identifiable ortholog at the expected location. A double dashed line indicates that synteny is not conserved but unable to be fully resolved with available data. White boxes with “n.d.” indicate that synteny was not examined, white boxes with “???” had no PCR amplification, and blank white boxes are spacers for visual display. Arrow directions reflect 5’ à 3’ orientation of the reading frame relative to the target gene (*e.g.*, <-Sas10-is oriented from 3’ to 5’ relative to –Msh5-> in the 5’ to 3’ direction.)

**Figure 3:**
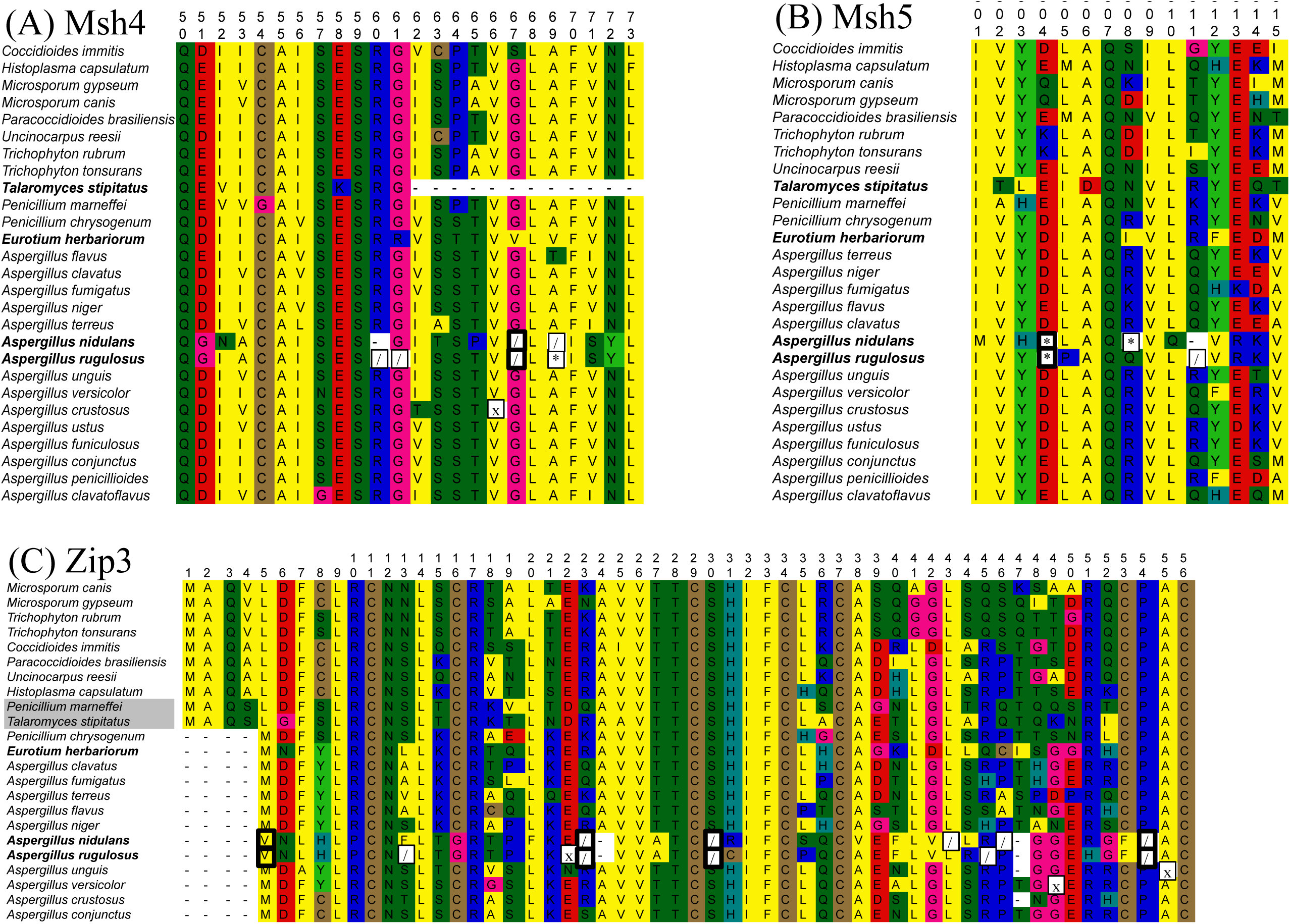
Candidate Non-Functionalizing Substitutions at Homologous Positions in *A. nidulans* and *A. rugulosus* Pseudogenes. Excerpts of (A) Msh4, (B) Msh5, and (C) Zip3 amino acid multiple sequence alignments; pseudogenized sequences have species names in bold, while possible incipient pseudogenes have species names highlighted in gray. Stop codons (box with *), frameshifts (box with /), and start codon alterations (black box around altered residue) at homologous positions have thicker box lines. The “x” boxes in *A. crustosus* Msh4, *A. unguis* Zip3, *A. versicolor* Zip3, and *A. rugulosus* Zip3 represent ambiguous residues due to base call ambiguity in the first position of each codon. (See supplemental file S1 for accession numbers; amino acid multiple sequence alignments without manual trimming will be in supplemental files S2-S5 (available on request in the interim on BioRxiv.)

**Figure 4:**
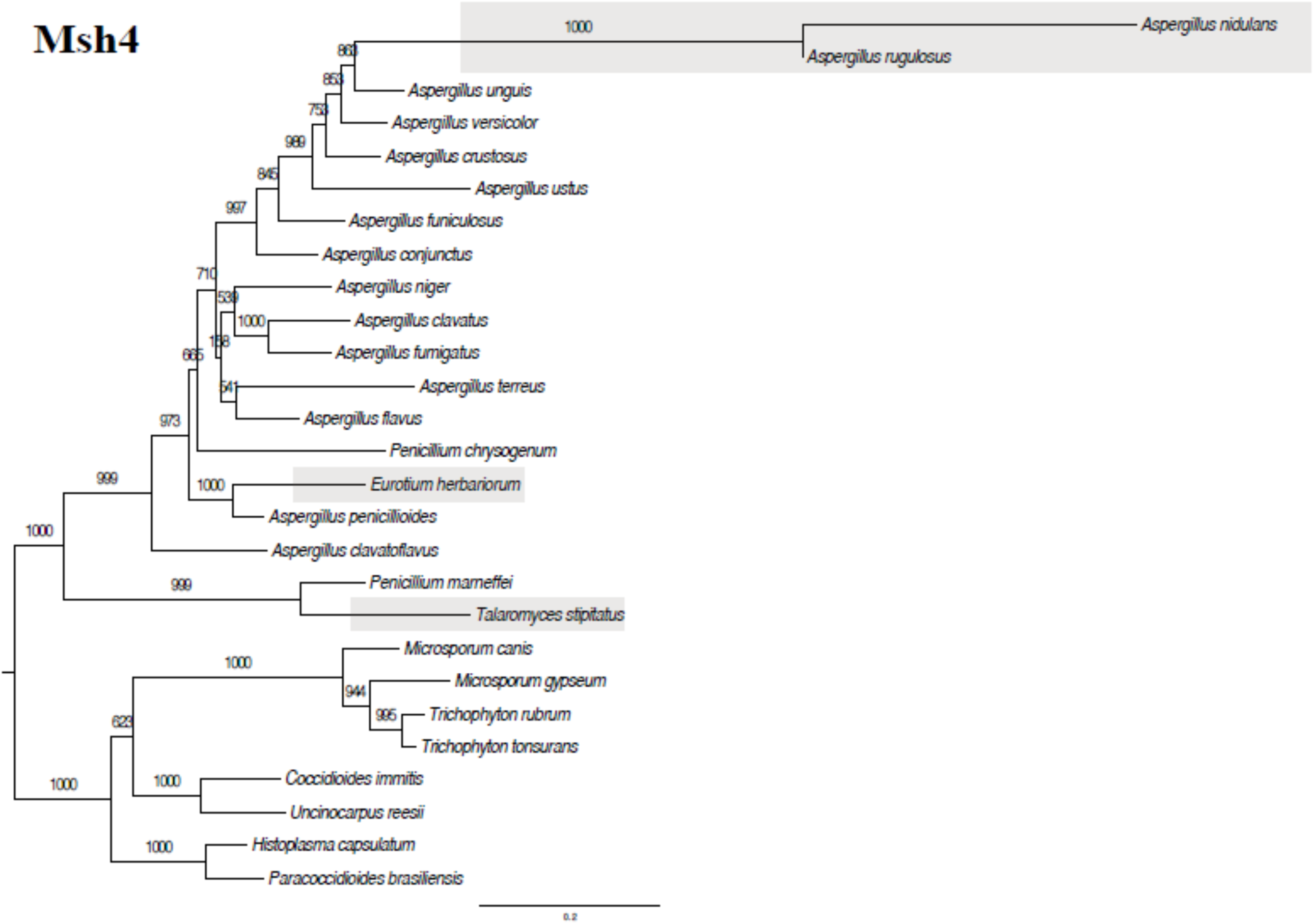
Maximum Likelihood Phylogeny of Msh4 Amino Acid Sequences. The Msh4 phylogeny was constructed from a 793-residue amino acid multiple sequence alignment in PhyML v3.0 (Guindon *et al*., 2010) using JTT+G5 with NNI and SPR search options and 1000 bootstrap replicates. Bootstrap support values are listed on each branch. Pseudogenes are denoted by a gray box.

**Figure 5:**
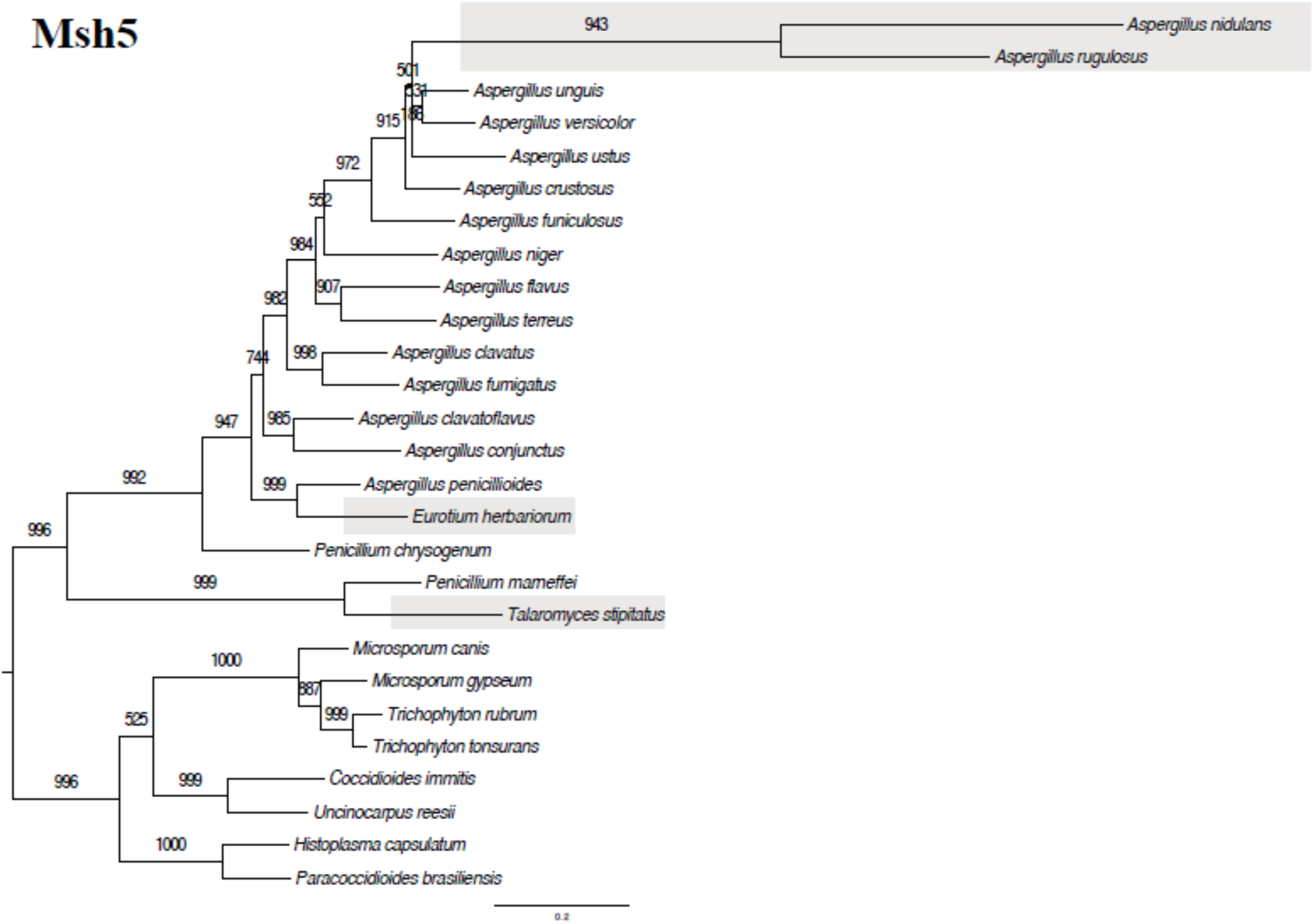
Maximum Likelihood Phylogeny of Msh5 Amino Acid Sequences. The Msh5 phylogeny was constructed from a 786-residue amino acid multiple sequence alignment in PhyML v3.0 (Guindon *et al*., 2010) using JTT+G5 with NNI and SPR search options and 1000 bootstrap replicates. Bootstrap support values are listed on each branch. Pseudogenes are denoted by a gray box.

**Table 1:** Comparison of Intact and Pseudogenized Msh4, Msh5, and Zip3

|  | Species | Status | Length<br>(bp<sup>[1]</sup>) | # Shifts | # Stops | # Splice Site<br>Disruptions | Length<br>(A.A.) | E vs. *A. unguis*<br>ortholog<sup>[2]</sup> |
| --- | --- | --- | --- | --- | --- | --- | --- | --- |
| **Msh4** | *A. unguis* | Intact | 3028 | 0 | 0 | 0 | 917 | n/a |
|  | *A. nidulans* | Pseudogene | 246 | 4 | 1 | 0 | 82 | 0.048 |
|  | *A. rugulosus* | Pseudogene | 1923 | 26 | 10 | 4 | 576 | 5E-60 |
|  | *E. herbariorum* | Pseudogene | 2963 | 2 | 3 | 0 | 892 | 0.0 |
|  | *T. stipitatus* | Pseudogene | 741 | 2 | 0 | 1 | 151 | 2E-25 |
| **Msh5** | *A. unguis* | Intact | 3682 | 0 | 0 | 1?<sup>[3]</sup> | 935 | n/a |
|  | *A. nidulans* | Pseudogene | 288 | 0 | 2 | 1 | 43 | 0.009 |
|  | *A. rugulosus* | Pseudogene | 704 | 3 | 3 | 5 | 169 | 4E-15 |
|  | *E. herbariorum* | Pseudogene | 3667 | 6 | 2 | 3 | 928 | 0.0 |
|  | *T. stipitatus* | Pseudogene | 2075 | 4 | 5 | 1 | 505 | 4E-53 |
| **Zip3** | *A. unguis* | Intact | 1291 | 0 | 0 | 0 | 362 | n/a |
|  | *A. nidulans* | Pseudogene | 1304 | 6 | 7 | 2 | 378 | 3E-30 |
|  | *A. rugulosus* | Pseudogene | 1168 | 14 | 4 | 2 | 274 | 1E-58 |
|  | *E. herbariorum* | Early Pseudogenization? | 1396 | 0 | 0 | 1 | 376 | 1E-81 |
|  | *T. stipitatus* | Pre-pseudogenization? | 1178 | 0 | 0 | 0 | 311 | 6E-34 |
|  | *P. marneffei* | Pre-pseudogenization? | 1248 | 0 | 0 | 0 | 338 | 3E-66 |

[1] Length includes introns and "N" placeholders for putative frameshifts
[2] BLASTX2SEQ vs. *A. unguis* protein prediction
[3] 1X local coverage

Eurotiomycete orthologs of Zip3 previously had been either only sporadically reported without phylogenetic validation (Chelysheva *et al*., 2012) or sought without success (Desjardins *et al*., 2011), so PSI-BLAST was used to identify a putative exemplar Eurotiomycete Zip3 ortholog in the NCBI nr protein database (*Trichophyton rubrum* hypothetical protein XP_003231222). This *T. rubrum* predicted protein sequence successfully served as a TBLASTN query to identify putative Zip3 orthologs in most of the other examined Eurotiomycetes (Figure 1, Figure 6), including *Penicillium marneffei* (with the manual annotation differing by only one intron from the XP_002145282 sequence identified by Chelysheva *et al*. [2012]). Similar to the other examined meiosis genes, TBLASTN searches did not identify a Zip3 ortholog in *A. nidulans* under parameters sufficient to retrieve the ortholog in congeners. Zip3 synteny was only moderately conserved across Eurotiomycetes: the 5’ flanking gene was usually a XAP5 domain protein (“XAP5”), but the 3’ flanking gene frequently varied from the consensus ATP citrate lyase subunit Acl (Figure 2). Nevertheless, the XAP5-Zip3-Acl synteny observed in many examined *Aspergillus* species allowed identification of the flanking gene orthologs and degrading Zip3 pseudogene in *A. nidulans* (Figure 2, Figure 3, Figure 6, Table 1).

**Figure 6:**
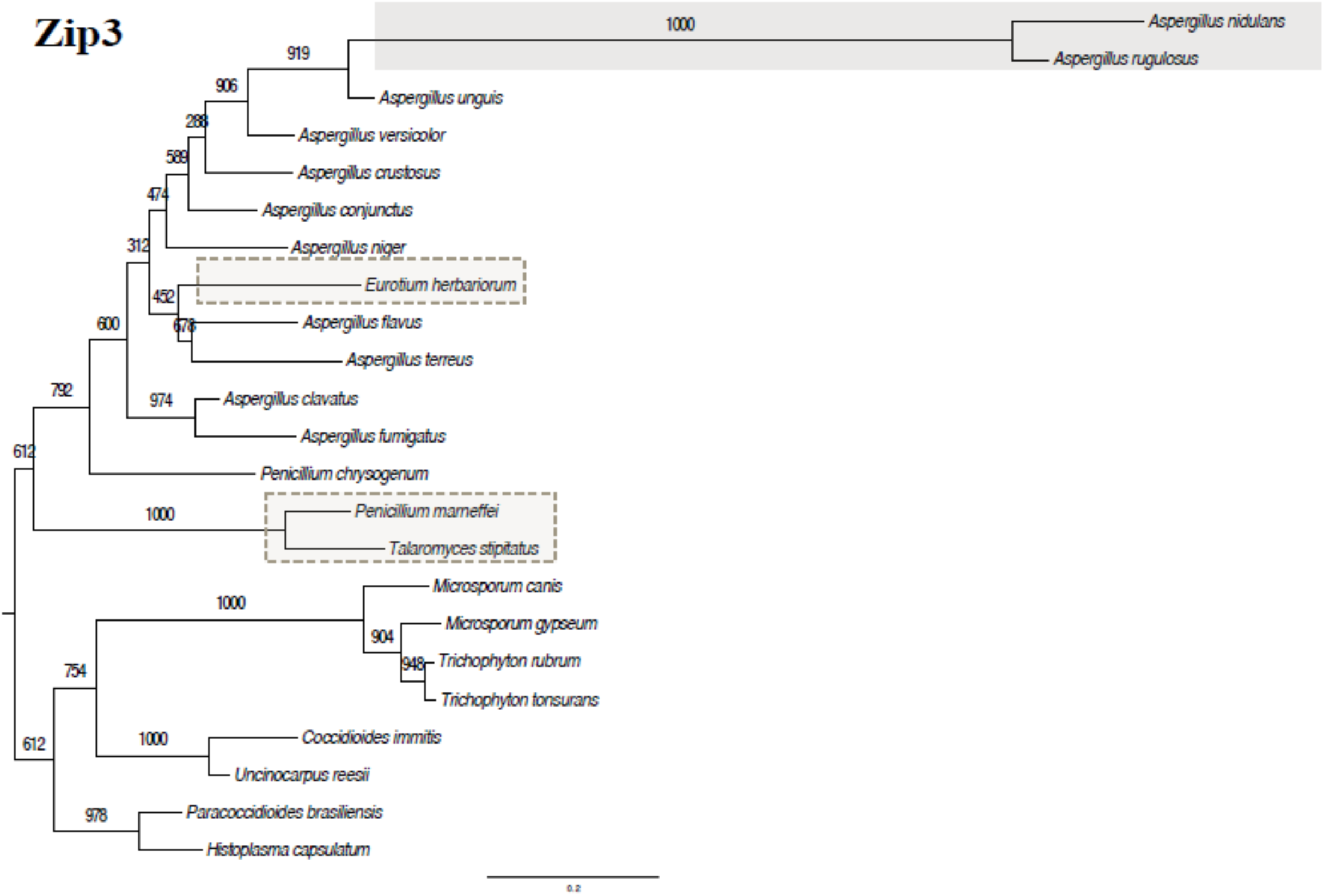
Maximum Likelihood Phylogeny of Zip3 Amino Acid Sequences. The Zip3 phylogeny was constructed from a 236-residue amino acid multiple sequence alignment in PhyML v3.0 (Guindon *et al*., 2010) using JTT+G5 with NNI and SPR search options and 1000 bootstrap replicates. Bootstrap support values are listed on each branch. Unambiguous pseudogenes are denoted by a gray box. Ambiguous, possibly pseudogenizing sequences are denoted by a dashed-line light gray box.

### Specific-Primer PCR Validation of the A. nidulans Genome

An initial confirmation step was to amplify and sequence the *A. nidulans* strain A4 flanking genes and intergenic spaces where Msh4, Msh5, and Pch2 are typically located under conserved Eurotiomycete synteny. The specific-primer and generate PCR amplification yielded sequences identical to those in the online genome assembly of *A. nidulans* strain A4 (Broad, 2013, also accessed 2012; data not shown), indicating that genome sequence misassembly was not the reason for failure to detect intact meiosis gene orthologs at the synteny-consensus locations.

### Pseudogene and Loss Distribution from Degenerate PCR and Additional Genome Sequences

The coinciding or consecutive nature of the four *A. nidulans* meiosis gene losses was investigated by performing degenerate PCR on nine additional *Aspergillus* species that lacked publicly available genome sequences at the time (*Aspergillus rugulosus, Aspergillus unguis, Aspergillus versicolor, Aspergillus crustosus, Aspergillus ustus, Aspergillus funiculosus, Aspergillus conjunctus, Aspergillus penicillioides, Aspergillus clavatoflavus*) using primers targeted to the conserved flanking genes (Msh4: Rsa4 and C2H2, Msh5: Hsp78 and Sas10, Pch2: Rcy1 and Cyt1, Zip3: XAP5 and Acl).

Eight of these nine additional *Aspergillus* species had intact Msh4 and Msh5 amplified; the ninth, *A. rugulosus*, contained degrading pseudogenes of Msh4 and Msh5 (Figure 1, Table 1, Figure 4, Figure 5; see supplemental file S1 for accession numbers) that each contained at least one possible nonfunctionalizing mutation at a position homologous to those in *A. nidulans* (Figure 3). The Eurotiomycete consensus synteny was recapitulated for the Msh4- and Msh5-flanking genes in all but one case (Figure 2): *A. penicillioides* Msh4 and C2H2 are separated by an inserted alpha-galactosidase (best NCBI BLASTX hit: *Microsporum canis* alpha-galactosidase A, EEQ28074, E-72.) The expected Zip3-containing region was not amplified from *A. ustus, A. funiculosus, A. penicillioides,* or *A. clavatoflavus* (Figure 1), which could be related to synteny around Zip3 being less conserved among the other examined Eurotiomycetes (Figure 2). Nevertheless, sequences that were amplified showed that Zip3 was pseudogenized in the same species in which Msh4 and Msh5 were pseudogenized (*A. rugulosus*) and intact in the other amplified sequences (Figure 1, Figure 6, Table 1; see supplemental file S1 for accession numbers). Also similar to Msh4 and Msh5, the *A. rugulosus* Zip3 pseudogene shared several potential nonfunctionalizing mutations with the *A. nidulans* Zip3 pseudogene (Figure 3).

Only a single putatively intact Pch2 ortholog was identified among the degenerate PCR taxa (Figure 1, Figure 7)—ironically, in a taxon with only partially conserved flanking gene synteny. The *A. penicillioides* Pch2 ortholog was obtained from a sequence amplified from expected primer binding at Cyt1 and serendipitous primer binding at the 5’ end of Pch2 itself. A partial Rcy1 sequence was amplified in *A. penicillioides*, so species-specific primers were used to try amplify the sequence between Rcy1 and the Pch2-Cyt1 region. The species-specific primers failed to yield amplification, which could be due to inversion, translocation, or large insertions on the 5’ side of *A. penicillioides* Pch2 (Figure 2). While *A. penicillioides* retains Pch2 despite its flanking gene synteny change, the remaining eight species retained flanking gene synteny but not Pch2 (Figure 1, Figure 2). These eight species with successful PCR amplification of sequences had positive BLAST identification of the flanking gene segments (data not shown) and substantial sequence similarity to the 5’ regulatory regions of Rcy1 and Cyt1 (Figure 8). However, no identifiable Pch2 sequence was found by manual inspection or BLASTX2SEQ comparisons to intact orthologs from *A. penicillioides, A. clavatus*, or *U. reesii*) (data not shown).

**Figure 7:**
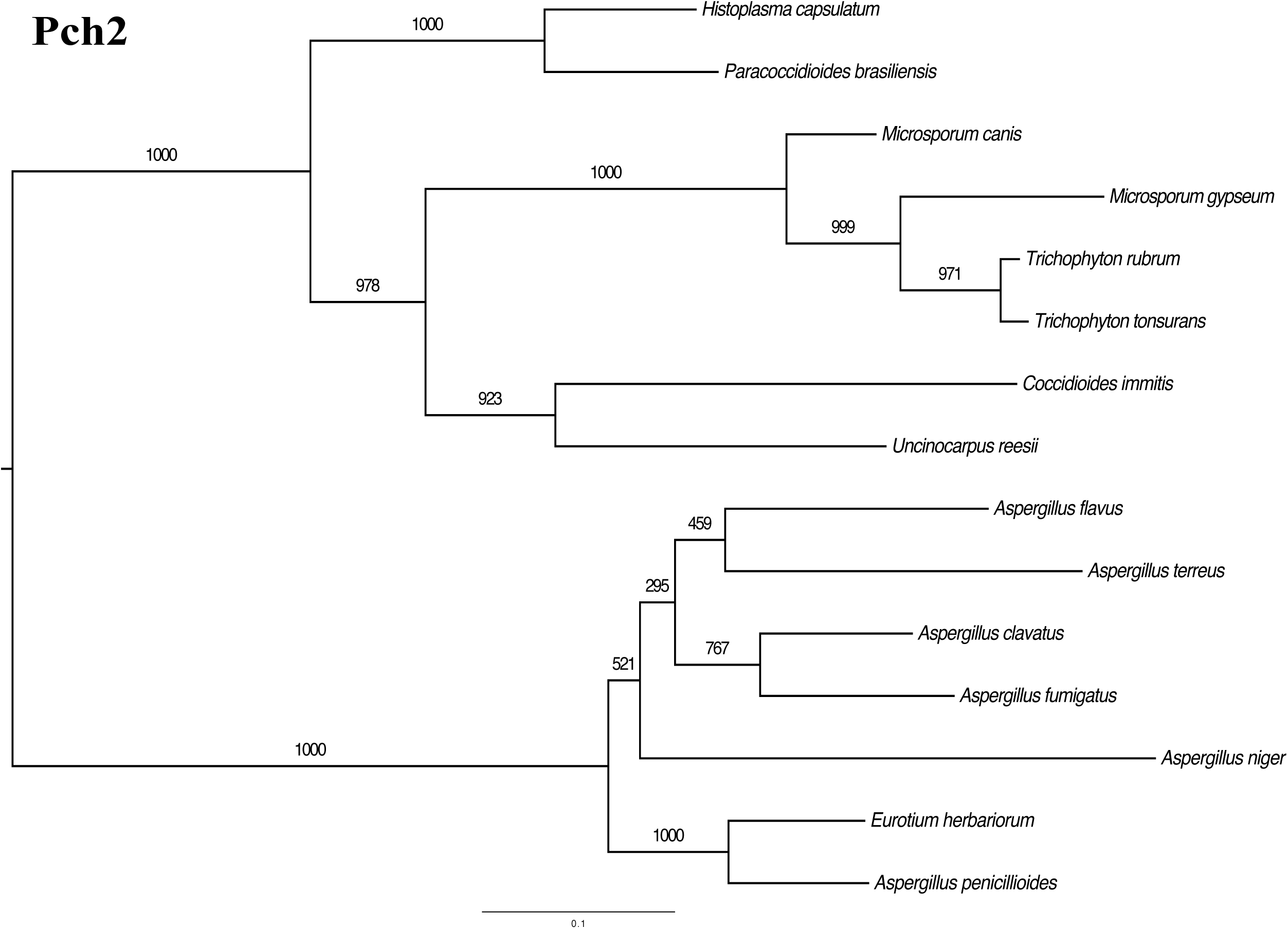
Maximum Likelihood Phylogeny of Pch2 Amino Acid Sequences. The Pch2 phylogeny was constructed from a 414-residue amino acid multiple sequence alignment in PhyML v3.0 (Guindon *et al*., 2010) using JTT+G5 with NNI and SPR search options and 1000 bootstrap replicates. Bootstrap support values are listed on each branch.

**Figure 8:**
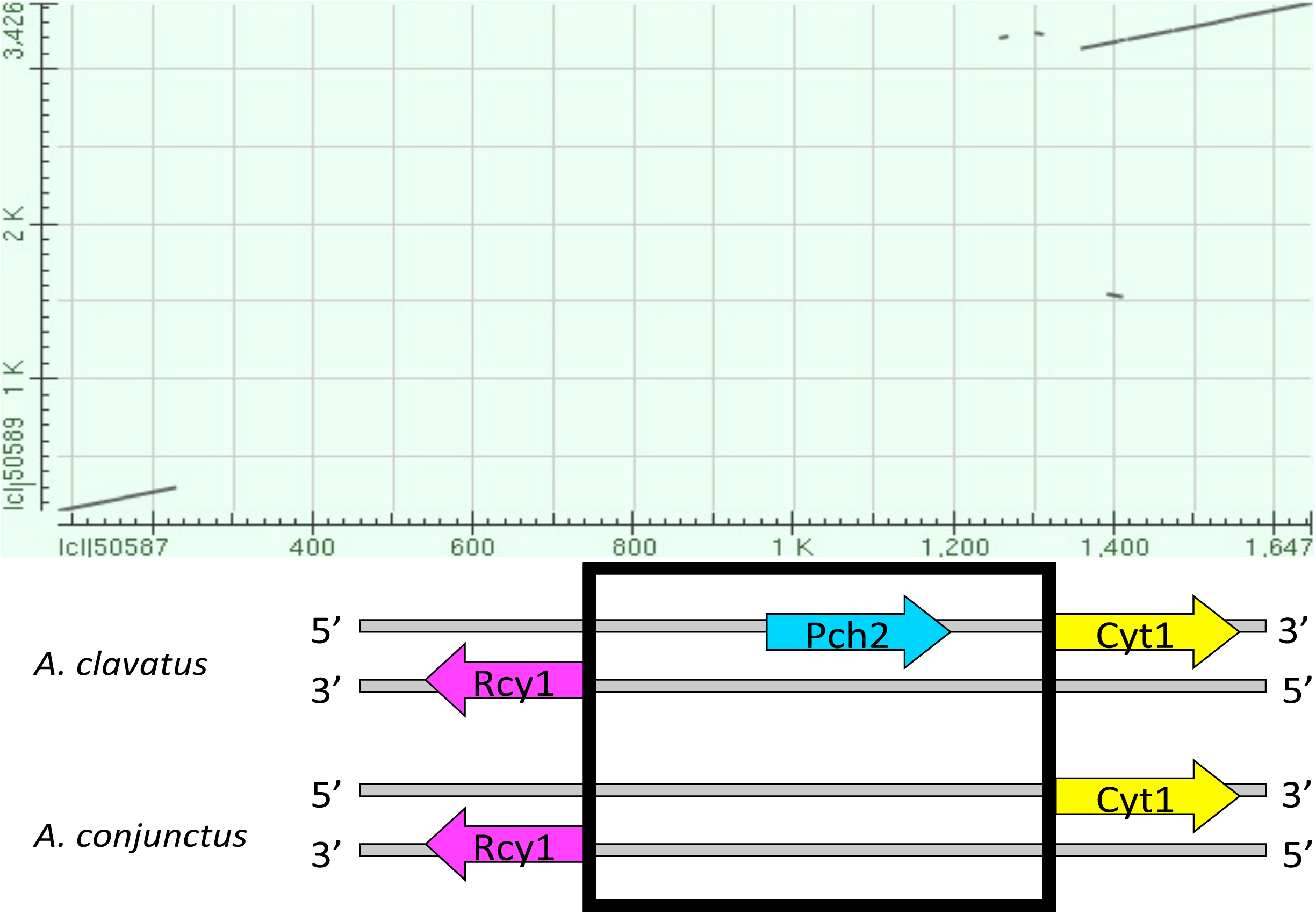
Dot Plot of Exemplar Rcy1-Pch2-Cyt1 and Rcy1-Cyt1 DNA. NCBI BLASTN2SEQ (Altschul *et al*., 1997; NCBI, 2012-2013) output is shown for the region between Rcy1 and Cyt1 in *A. clavatus* (exemplar species with intact Pch2) and *A. conjunctus* (exemplar species lacking intact Pch2), with the aligned region marked by the red box in the synteny diagram. Dark gray graph markings show significant nucleotide sequence alignment at the positions noted on the axes (X: *A. conjunctus*, Y: *A. clavatus*.)

To test whether the lack of Pch2 at the expected syntenic location in the degenerate PCR work corresponded to an inability to detect Pch2 elsewhere in the genome of related taxa, additional Eurotiomycete genome sequences (genome information in supplemental file S1) were examined with TBLASTN, BLASTX, and manual annotation (Figure 1 taxa with “g” notation). *Aspergillus versicolor*, *Aspergillus sydowii*, and *Aspergillus zonatus* each had no detected ortholog of Pch2 (Figure 1) despite the parameters being sufficient to return numerous nonorthologous AAA ATPases (data not shown). Importantly, the *A. versicolor* genome sequence BLAST results independently replicate the degenerate PCR results (with Pch2 undetected by both methods) with a similar lack of detected Pch2 in this species’ close relative *A. sydowii* (Peterson, 2008). *A. zonatus* is a sister taxon to *A. clavatoflavus* among the examined Eurotiomycetes (Peterson, 2008), which could suggest that Pch2 was lost in the most recent common ancestor of those two species. Together, these analyses and the original Pch2 searches in *A. nidulans* are suggestive of Pch2 loss at the syntenic locus equating to loss of Pch2 from the genome. Msh4, Msh5, and Zip3 were also present and intact in *A. sydowii* and *A. zonatus* (Figure 1, supplemental file S1). All four genes of interest were present and apparently intact in the remaining taxa examined by BLAST searches and manual annotation only (*A. acidus, A. aculeatus, A. brasiliensis, A. carbonarius, A. glaucus, A. oryzae, A. tubingensis, A. wentii, Arthroderma benhamiae, Coccidioides posadasii, Neosartorya fischeri, Trichophyton equinum, Trichophyton verrucosum*, Figure 1).

### Topology Tests

The relationship of *A. penicillioides* and *A. clavatoflavus* relative to the other examined *Aspergillus* clades was not well-resolved in other studies (Peterson, 2008). Determining this relationship was necessary to (i) infer the number of independent Pch2 losses in genus *Aspergillus* and (ii) have a defined reference topology for K_a_/K_s_ comparisons. Topology comparison tests in TREE-PUZZLE (Schmidt *et al*., 2002) found an arrangement with *A. clavatoflavus* most basal, *A. penicillioides* second most basal, and *A. nidulans-A.conjunctus* sister to *A. clavatus*-*A. niger* (“topology 1” in Figure 9) to be the only tested topology to never be rejected as significantly worse than the best topology (which also was “topology 1” itself in the analyses using either Msh4 or concatenated Msh4 and Msh5, Figure 9). Therefore, topology 1 was utilized in subsequent Codeml tests. Notably, both tested topologies representing a single loss of Pch2 (*A. penicillioides* sister to *A. nidulans-A. conjunctus*, topologies 6 and 10 in Figure 9) were rejected by all four statistical tests using Msh4 and concatenated Msh4 and Msh5 alignments; although these “single-loss scenario” topologies could not be rejected as significantly worse than the best tested tree using Msh5, use of the Msh5 sequences led to none of the tested topologies (single-loss or multiple-loss) being rejected at all.

**Figure 9:**
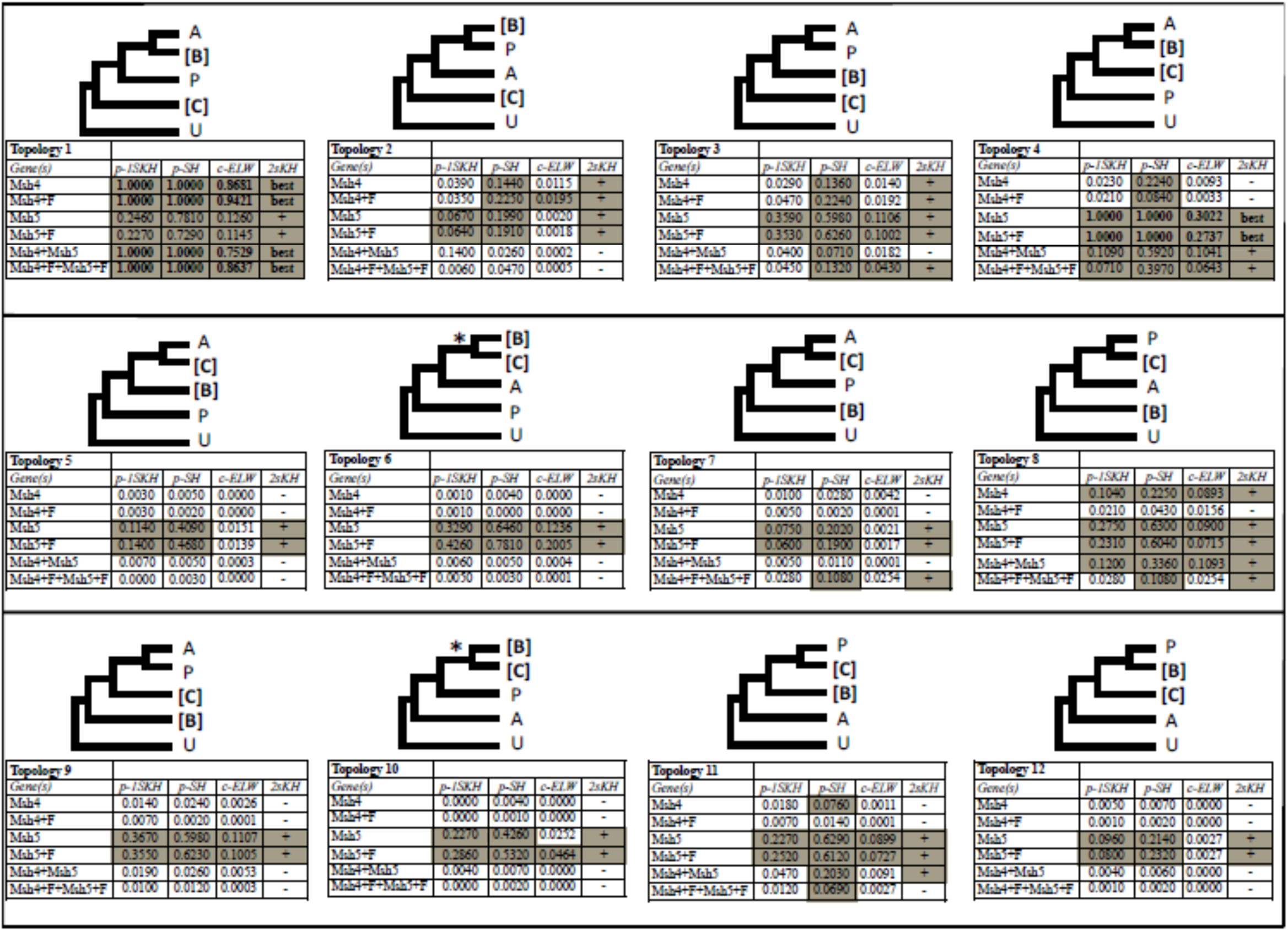
TREE-PUZZLE Topology Test Comparisons. Letters A and B represent well-supported clades in Peterson (2008) (A: *A. terreus, A. flavus, A. niger, A. clavatus, A. fumigatus, N. fischeri*; B: *A. nidulans, A. rugulosus, A. unguis, A. versicolor, A. crustosus, A. ustus, A. funiculosus, A. conjunctus*). Relationships within clade A and clade B followed Peterson (2008) (see Figure 1.) Letter P represents *A. penicillioides*, letter C represents *A. clavatoflavus*, and letter U represents *Uncinocarpus reesii* (outgroup). Bold brackets around B and C indicate a derived loss of Pch2, and the number of independent Pch2 losses is stated below the topology.

### Comparison of K_a_/K_s_ in Msh4, Msh5, and Zip3 Relative to Pch2 Losses

To determine whether the inferred loss of Pch2 in the most recent common ancestor (MRCA) of *A. nidulans* and *A. conjunctus* was associated with changes in the sequence evolution patterns of Msh4, Msh5, and/or Zip3 prior to their later pseudogenization in the MRCA of *A. nidulans* and *A. rugulosus*, K_a_/K_s_ comparisons were conducted in Codeml (Yang, 2007). Saturation (K_s_ > 1) was evident for some terminal branches in the Msh4 (*A. terreus, A. niger, A. clavatoflavus*) and Msh5 (*A. terreus, A. niger, A. conjunctus, A. penicillioides*) analyses and the *U. reesii* outgroup branch, but most branches had K_s_ < 1.

Despite varying the inclusion or exclusion of pseudogenes (Table 2A), assignment of a separate K_a_/K_s_ ratio to pseudogenes (Table 2B), including or excluding the independent *A. clavatoflavus* Pch2 loss (Table 2B *vs.* Table 2C-2D), and assigning a separate K_a_/K_s_ ratio to varied subsets of Pch2-lacking species with intact ZMM genes (Table 2B-2D), no evidence was found for a pervasive elevation or depression of Msh4 K_a_/K_s_ in Pch2-lacking *Aspergillus*. When assumed to have an underlying K_a_/K_s_ ratio separate from all other intact *Aspergillus* Msh4 sequences, *A. ustus* Msh4 did have a modestly but significantly higher K_a_/K_s_ than other intact *Aspergillus* Msh4 (0.0959 *vs.* 0.0498, p<0.001, Table 2D). The relatively higher K_a_/K_s_ of *A. ustus* Msh4 is likely relevant to the results for some two-ratio models (one ratio for a subset of Pch2-lacking taxa, one ratio for all remaining taxa, pseudogenes excluded; Table 2D): the two-ratio models were weakly significantly better than the null one-ratio model (0.01<p<0.05) only when *A. ustus* Msh4 and a few additional species were included in the “taxon subset” ratio category (Table 2D). This suggests that, while *A. ustus* has an apparently elevated Msh4 K_a_/K_s_, there is no evidence for systematic difference among the other intact *Aspergillus* Msh4 sequences whether Pch2 is intact or not. (The Msh4 pseudogenes, as expected, were supported as having a substantially higher underlying K_a_/K_s_ than the intact orthologs (Table 2A-2B.))

**Table 2:**
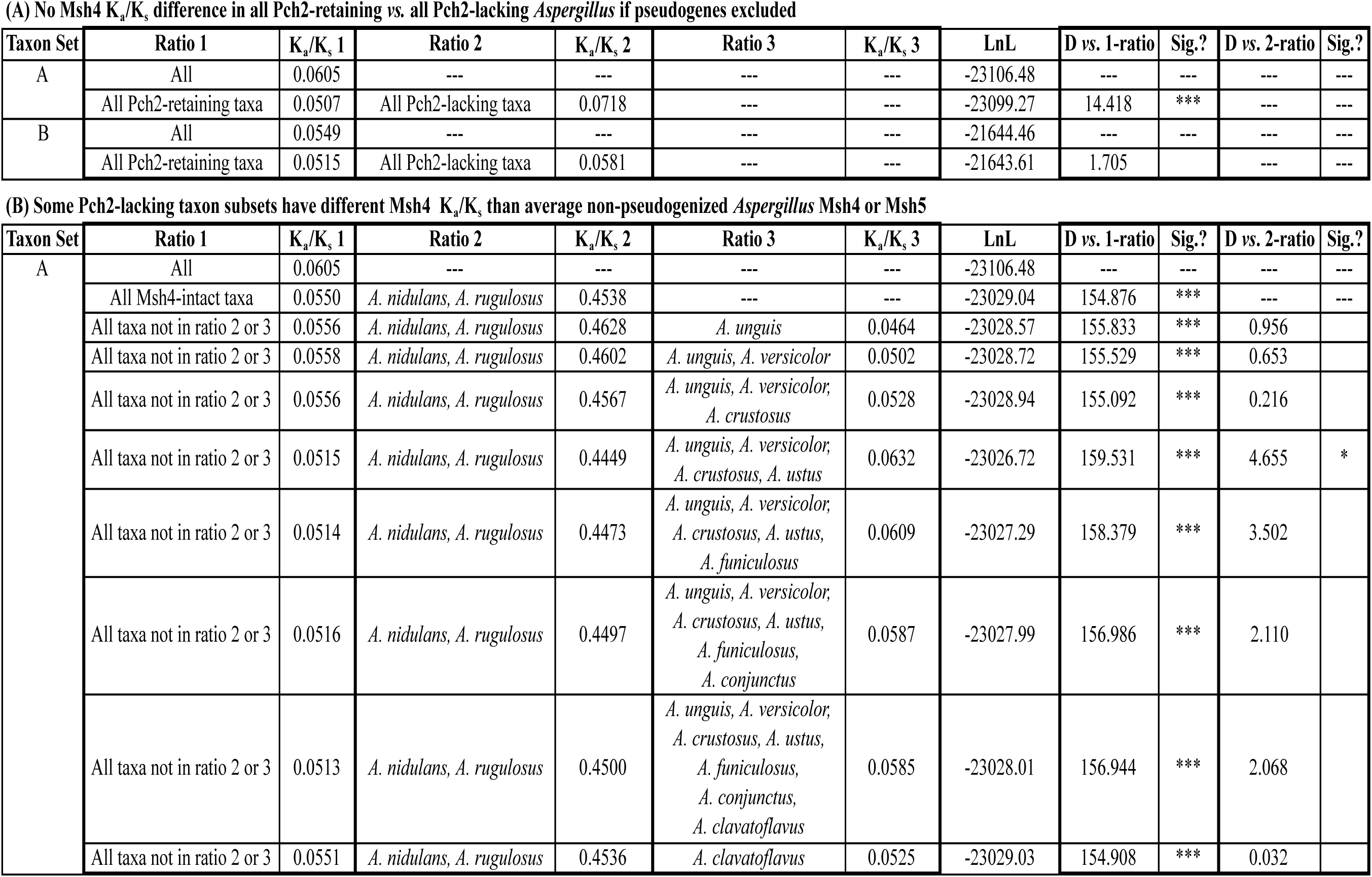
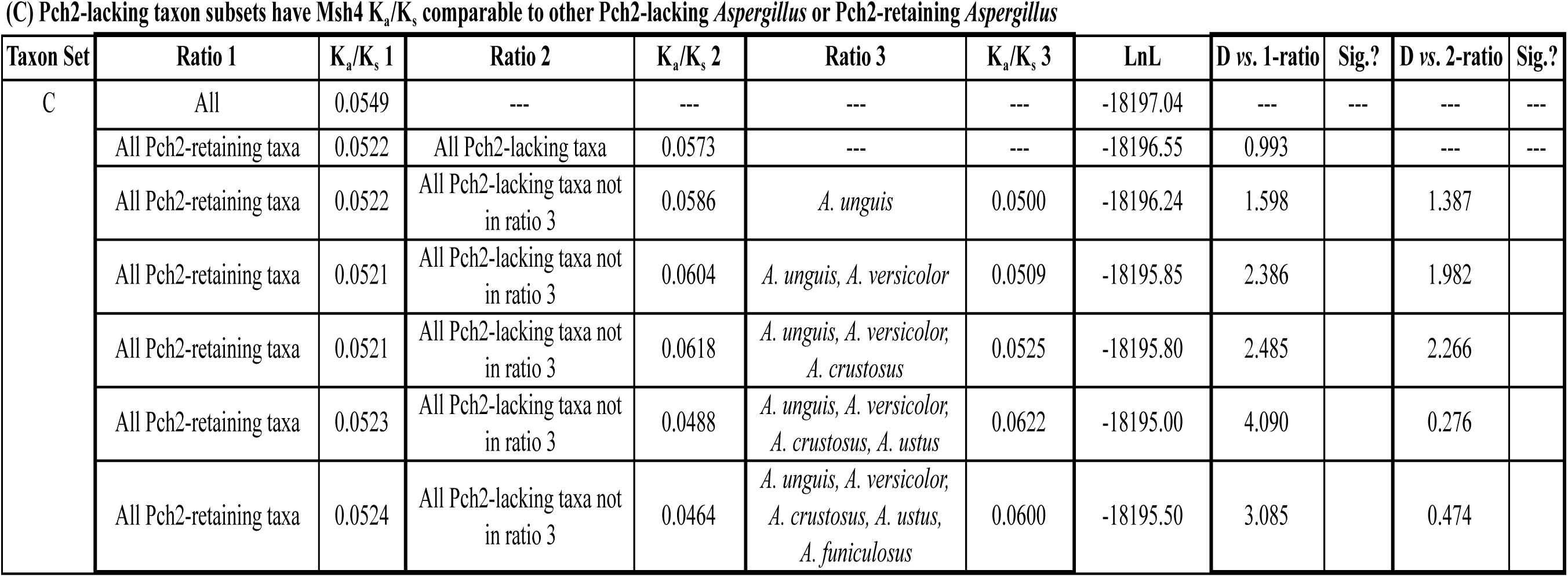
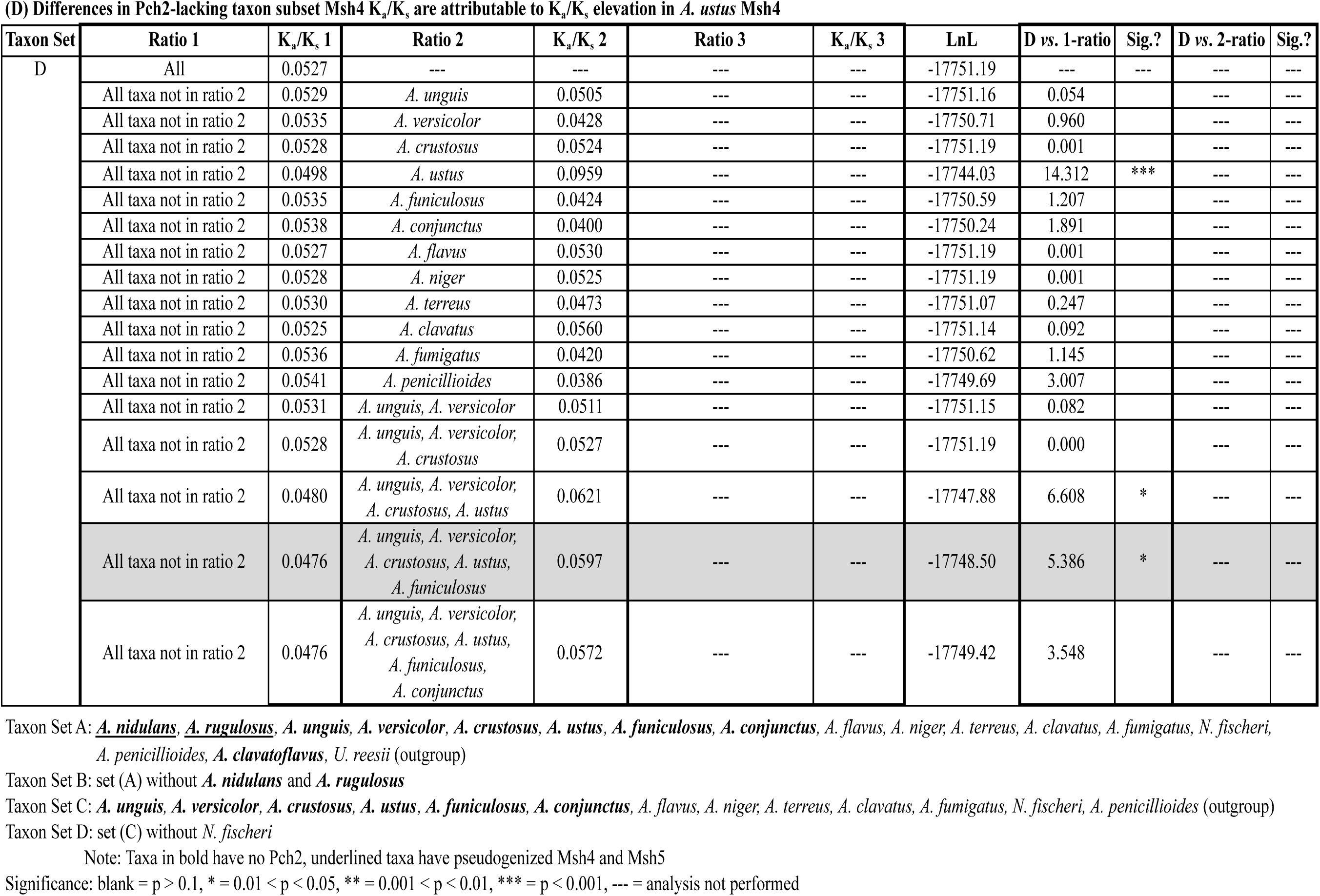
Codeml Analysis of Msh4 With Respect to Pch2 Loss.

**(A) No Msh4 $K_a/K_s$ difference in all Pch2-retaining vs. all Pch2-lacking *Aspergillus* if pseudogenes excluded**
| Taxon Set | Ratio 1 | $K_a/K_s$ 1 | Ratio 2 | $K_a/K_s$ 2 | Ratio 3 | $K_a/K_s$ 3 | LnL | D vs. 1-ratio | Sig.? | D vs. 2-ratio | Sig.? |
| --- | --- | --- | --- | --- | --- | --- | --- | --- | --- | --- | --- |
| A | All | 0.0605 | --- | --- | --- | --- | -23106.48 | --- | --- | --- | --- |
|  | All Pch2-retaining taxa | 0.0507 | All Pch2-lacking taxa | 0.0718 | --- | --- | -23099.27 | 14.418 | *** | --- | --- |
| B | All | 0.0549 | --- | --- | --- | --- | -21644.46 | --- | --- | --- | --- |
|  | All Pch2-retaining taxa | 0.0515 | All Pch2-lacking taxa | 0.0581 | --- | --- | -21643.61 | 1.705 |  | --- | --- |

| Taxon Set | Ratio 1 | $K_a/K_s$ 1 | Ratio 2 | $K_a/K_s$ 2 | Ratio 3 | $K_a/K_s$ 3 | LnL | D vs. 1-ratio | Sig.? | D vs. 2-ratio | Sig.? |
| --- | --- | --- | --- | --- | --- | --- | --- | --- | --- | --- | --- |
| A | All | 0.0605 | --- | --- | --- | --- | -23106.48 | --- | --- | --- | --- |
|  | All Msh4-intact taxa | 0.0550 | *A. nidulans*, *A. rugulosus* | 0.4538 | --- | --- | -23029.04 | 154.876 | *** | --- | --- |
|  | All taxa not in ratio 2 or 3 | 0.0556 | *A. nidulans*, *A. rugulosus* | 0.4628 | *A. unguis* | 0.0464 | -23028.57 | 155.833 | *** | 0.956 |  |
|  | All taxa not in ratio 2 or 3 | 0.0558 | *A. nidulans*, *A. rugulosus* | 0.4602 | *A. unguis*, *A. versicolor* | 0.0502 | -23028.72 | 155.529 | *** | 0.653 |  |
|  | All taxa not in ratio 2 or 3 | 0.0556 | *A. nidulans*, *A. rugulosus* | 0.4567 | *A. unguis*, *A. versicolor*,<br>*A. crustosus* | 0.0528 | -23028.94 | 155.092 | *** | 0.216 |  |
|  | All taxa not in ratio 2 or 3 | 0.0515 | *A. nidulans*, *A. rugulosus* | 0.4449 | *A. unguis*, *A. versicolor*,<br>*A. crustosus*, *A. ustus* | 0.0632 | -23026.72 | 159.531 | *** | 4.655 | * |
|  | All taxa not in ratio 2 or 3 | 0.0514 | *A. nidulans*, *A. rugulosus* | 0.4473 | *A. unguis*, *A. versicolor*,<br>*A. crustosus*, *A. ustus*,<br>*A. funiculosus* | 0.0609 | -23027.29 | 158.379 | *** | 3.502 |  |
|  | All taxa not in ratio 2 or 3 | 0.0516 | *A. nidulans*, *A. rugulosus* | 0.4497 | *A. unguis*, *A. versicolor*,<br>*A. crustosus*, *A. ustus*,<br>*A. funiculosus*,<br>*A. conjunctus* | 0.0587 | -23027.99 | 156.986 | *** | 2.110 |  |
|  | All taxa not in ratio 2 or 3 | 0.0513 | *A. nidulans*, *A. rugulosus* | 0.4500 | *A. unguis*, *A. versicolor*,<br>*A. crustosus*, *A. ustus*,<br>*A. funiculosus*,<br>*A. conjunctus*,<br>*A. clavatoflavus* | 0.0585 | -23028.01 | 156.944 | *** | 2.068 |  |
|  | All taxa not in ratio 2 or 3 | 0.0551 | *A. nidulans*, *A. rugulosus* | 0.4536 | *A. clavatoflavus* | 0.0525 | -23029.03 | 154.908 | *** | 0.032 |  |
| C | All | 0.0549 | --- | --- | --- | --- | -18197.04 | --- | --- | --- | --- |
|  | All Pch2-retaining taxa | 0.0522 | All Pch2-lacking taxa | 0.0573 | --- | --- | -18196.55 | 0.993 |  | --- | --- |
|  | All Pch2-retaining taxa | 0.0522 | All Pch2-lacking taxa not in ratio 3 | 0.0586 | *A. unguis* | 0.0500 | -18196.24 | 1.598 |  | 1.387 |  |
|  | All Pch2-retaining taxa | 0.0521 | All Pch2-lacking taxa not in ratio 3 | 0.0604 | *A. unguis*, *A. versicolor* | 0.0509 | -18195.85 | 2.386 |  | 1.982 |  |
|  | All Pch2-retaining taxa | 0.0521 | All Pch2-lacking taxa not in ratio 3 | 0.0618 | *A. unguis*, *A. versicolor*,<br>*A. crustosus* | 0.0525 | -18195.80 | 2.485 |  | 2.266 |  |
|  | All Pch2-retaining taxa | 0.0523 | All Pch2-lacking taxa not in ratio 3 | 0.0488 | *A. unguis*, *A. versicolor*,<br>*A. crustosus*, *A. ustus* | 0.0622 | -18195.00 | 4.090 |  | 0.276 |  |
|  | All Pch2-retaining taxa | 0.0524 | All Pch2-lacking taxa not in ratio 3 | 0.0464 | *A. unguis*, *A. versicolor*,<br>*A. crustosus*, *A. ustus*,<br>*A. funiculosus* | 0.0600 | -18195.50 | 3.085 |  | 0.474 |  |
| D | All | 0.0527 | --- | --- | --- | --- | -17751.19 | --- | --- | --- | --- |
|  | All taxa not in ratio 2 | 0.0529 | *A. unguis* | 0.0505 | --- | --- | -17751.16 | 0.054 |  | --- | --- |
|  | All taxa not in ratio 2 | 0.0535 | *A. versicolor* | 0.0428 | --- | --- | -17750.71 | 0.960 |  | --- | --- |
|  | All taxa not in ratio 2 | 0.0528 | *A. crustosus* | 0.0524 | --- | --- | -17751.19 | 0.001 |  | --- | --- |
|  | All taxa not in ratio 2 | 0.0498 | *A. ustus* | 0.0959 | --- | --- | -17744.03 | 14.312 | *** | --- | --- |
|  | All taxa not in ratio 2 | 0.0535 | *A. funiculosus* | 0.0424 | --- | --- | -17750.59 | 1.207 |  | --- | --- |
|  | All taxa not in ratio 2 | 0.0538 | *A. conjunctus* | 0.0400 | --- | --- | -17750.24 | 1.891 |  | --- | --- |
|  | All taxa not in ratio 2 | 0.0527 | *A. flavus* | 0.0530 | --- | --- | -17751.19 | 0.001 |  | --- | --- |
|  | All taxa not in ratio 2 | 0.0528 | *A. niger* | 0.0525 | --- | --- | -17751.19 | 0.001 |  | --- | --- |
|  | All taxa not in ratio 2 | 0.0530 | *A. terreus* | 0.0473 | --- | --- | -17751.07 | 0.247 |  | --- | --- |
|  | All taxa not in ratio 2 | 0.0525 | *A. clavatus* | 0.0560 | --- | --- | -17751.14 | 0.092 |  | --- | --- |
|  | All taxa not in ratio 2 | 0.0536 | *A. fumigatus* | 0.0420 | --- | --- | -17750.62 | 1.145 |  | --- | --- |
|  | All taxa not in ratio 2 | 0.0541 | *A. penicillioides* | 0.0386 | --- | --- | -17749.69 | 3.007 |  | --- | --- |
|  | All taxa not in ratio 2 | 0.0531 | *A. unguis, A. versicolor* | 0.0511 | --- | --- | -17751.15 | 0.082 |  | --- | --- |
|  | All taxa not in ratio 2 | 0.0528 | *A. unguis, A. versicolor, A. crustosus* | 0.0527 | --- | --- | -17751.19 | 0.000 |  | --- | --- |
|  | All taxa not in ratio 2 | 0.0480 | *A. unguis, A. versicolor, A. crustosus, A. ustus* | 0.0621 | --- | --- | -17747.88 | 6.608 | * | --- | --- |
|  | All taxa not in ratio 2 | 0.0476 | *A. unguis, A. versicolor, A. crustosus, A. ustus, A. funiculosus* | 0.0597 | --- | --- | -17748.50 | 5.386 | * | --- | --- |
|  | All taxa not in ratio 2 | 0.0476 | *A. unguis, A. versicolor, A. crustosus, A. ustus, A. funiculosus, A. conjunctus* | 0.0572 | --- | --- | -17749.42 | 3.548 |  | --- | --- |

Taxon Set A: *A. nidulans*, *A. rugulosus*, *A. unguis*, *A. versicolor*, *A. crustosus*, *A. ustus*, *A. funiculosus*, *A. conjunctus*, *A. flavus*, *A. niger*, *A. terreus*, *A. clavatus*, *A. fumigatus*, *N. fischeri*, *A. penicillioides*, *A. clavatoflavus*, *U. reesii* (outgroup)
Taxon Set B: set (A) without *A. nidulans* and *A. rugulosus*
Taxon Set C: *A. unguis*, *A. versicolor*, *A. crustosus*, *A. ustus*, *A. funiculosus*, *A. conjunctus*, *A. flavus*, *A. niger*, *A. terreus*, *A. clavatus*, *A. fumigatus*, *N. fischeri*, *A. penicillioides* (outgroup)
Taxon Set D: set (C) without *N. fischeri*
Note: Taxa in bold have no Pch2, underlined taxa have pseudogenized Msh4 and Msh5
Significance: blank = $p > 0.1$ , \* = $0.01 < p < 0.05$ , \*\* = $0.001 < p < 0.01$ , \*\*\* = $p < 0.001$ , --- = analysis not performed

The results for K_a_/K_s_ comparisons of *Aspergillus* Msh5 (Table 3) and Zip3 (Table 4) relative to Pch2 loss status were similar to the Msh4 results. The pseudogenized *A. nidulans* and *A. rugulosus* Msh5 and Zip3 sequences were consistently supported as having higher K_a_/K_s_ ratios than intact orthologs (two-ratio model *vs.* null one-ratio model in Table 3B, Table 4), but no consistent difference in intact Msh5 or Zip3 K_a_/K_s_ with respect to Pch2 loss was found. Some models assigning a different Msh5 K_a_/K_s_ ratio to subsets of Pch2-lacking *Aspergillus* were significantly better (p<0.05) than models assuming the same ratio for all intact Msh5 (Table 3B-3C), but those results are likely attributable to the atypically low K_a_/K_s_ of *A. ustus* Msh5 (0.0457 *vs*. 0.0722 in all other intact Msh5, 0.01<p<0.05, Table 3D) and the atypically high K_a_/K_s_ of *A. flavus* Msh5 (0.1060 *vs.* 0.0677 in all other intact Msh5, 0.01<p<0.05, Table 3D). No individual taxa or subsets of Pch2-lacking taxa had notably different Zip3 K_a_/K_s_ ratios. These findings suggest there was not a general relaxation of sequence evolution constraints on Msh4, Msh5, or Zip3 coincident with loss of Pch2 in the *A. nidulans-A. conjunctus* clade. Furthermore, there was not a concordant increase in K_a_/K_s_ of Msh4, Msh5, and Zip3 together in any of the examined Pch2-lacking *Aspergillus* that diverged prior to the speciation of *A. nidulans* and *A. rugulosus* (*i.e.*, only the taxa in which Msh4, Msh5, and Zip3 pseudogenized appear to have had K_a_/K_s_ elevations in all three genes—no preceding modest partial relaxation of sequence evolution constraints in species like *A. unguis* or *A. versicolor* was observed.)

**Table 3:**
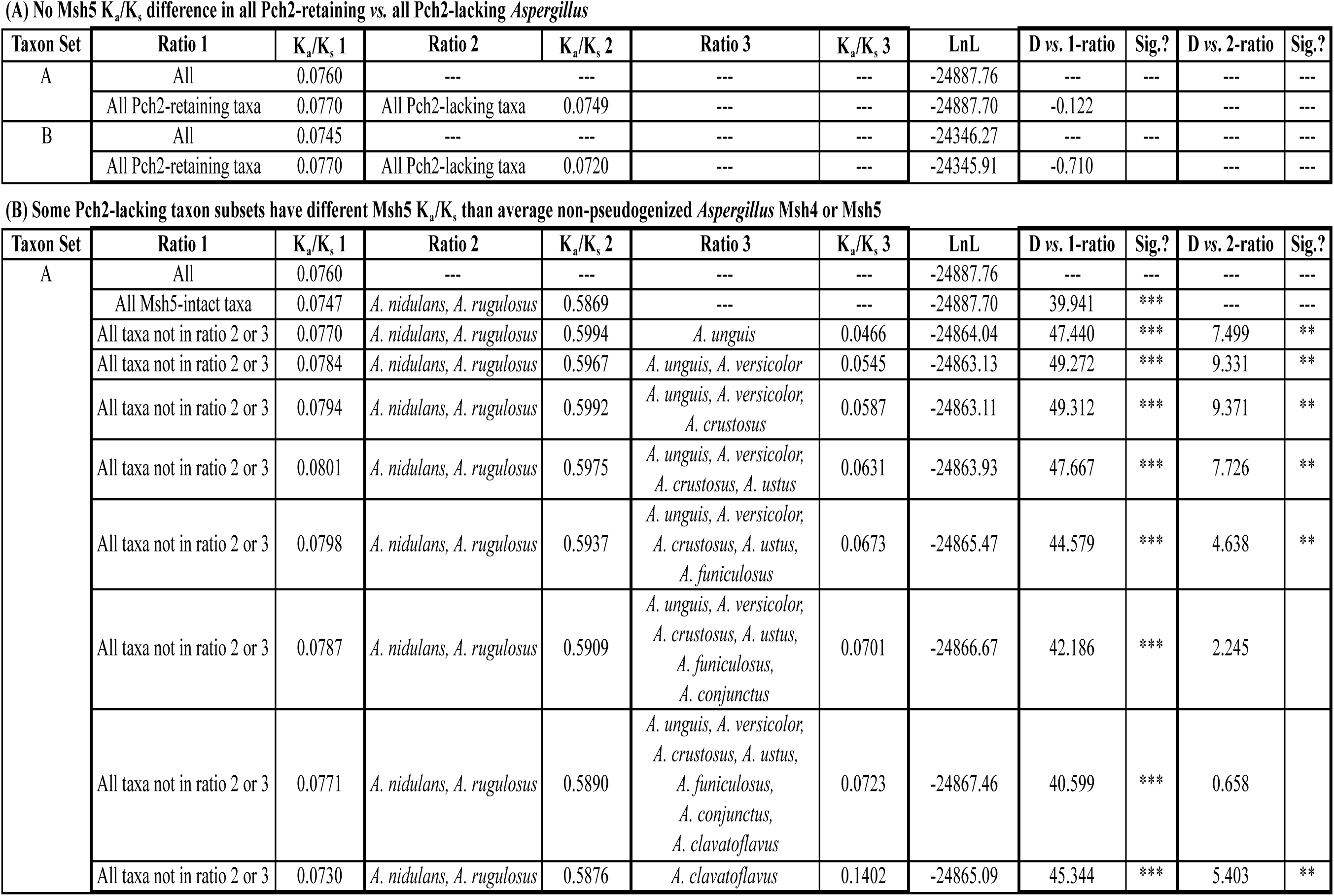
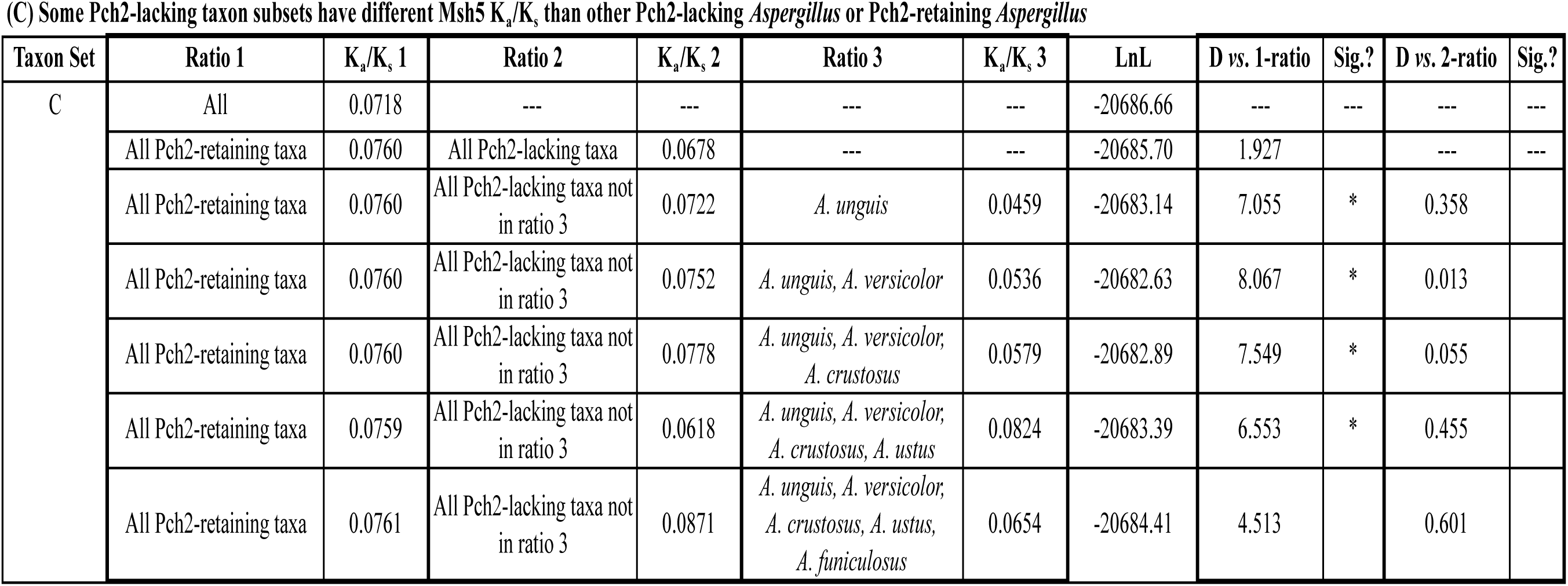
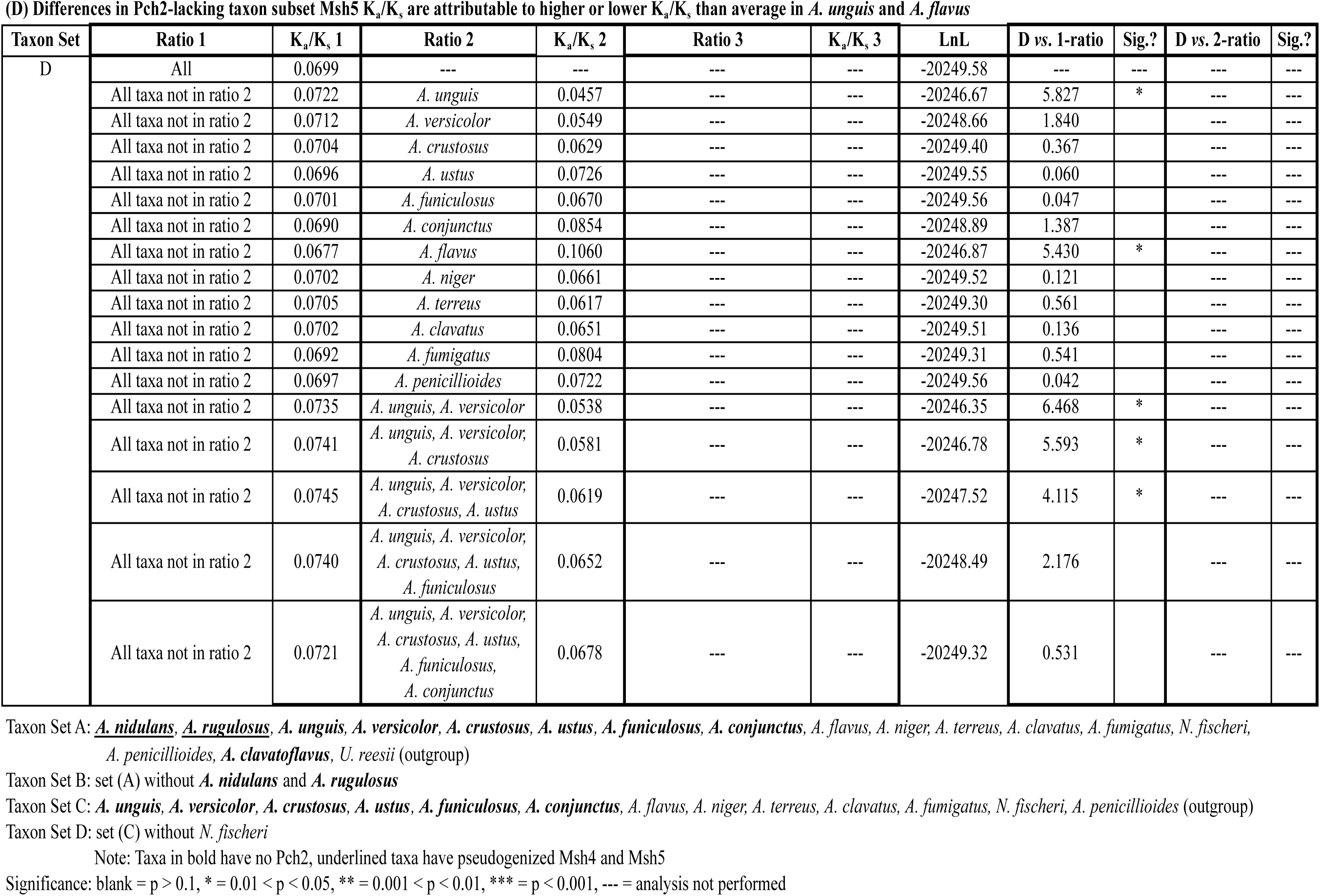
Codeml Analysis of Msh5 With Respect to Pch2 Loss.

No Msh5 $K_a/K_s$ difference in all Pch2-retaining vs. all Pch2-lacking *Aspergillus*
| Taxon Set | Ratio 1 | $K_a/K_s$ 1 | Ratio 2 | $K_a/K_s$ 2 | Ratio 3 | $K_a/K_s$ 3 | LnL | D vs. 1-ratio | Sig.? | D vs. 2-ratio | Sig.? |
| --- | --- | --- | --- | --- | --- | --- | --- | --- | --- | --- | --- |
| A | All | 0.0760 | --- | --- | --- | --- | -24887.76 | --- | --- | --- | --- |
|  | All Pch2-retaining taxa | 0.0770 | All Pch2-lacking taxa | 0.0749 | --- | --- | -24887.70 | -0.122 |  | --- | --- |
| B | All | 0.0745 | --- | --- | --- | --- | -24346.27 | --- | --- | --- | --- |
|  | All Pch2-retaining taxa | 0.0770 | All Pch2-lacking taxa | 0.0720 | --- | --- | -24345.91 | -0.710 |  | --- | --- |

| Taxon Set | Ratio 1 | $K_a/K_s$ 1 | Ratio 2 | $K_a/K_s$ 2 | Ratio 3 | $K_a/K_s$ 3 | LnL | D vs. 1-ratio | Sig.? | D vs. 2-ratio | Sig.? |
| --- | --- | --- | --- | --- | --- | --- | --- | --- | --- | --- | --- |
| A | All | 0.0760 | --- | --- | --- | --- | -24887.76 | --- | --- | --- | --- |
|  | All Msh5-intact taxa | 0.0747 | *A. nidulans*, *A. rugulosus* | 0.5869 | --- | --- | -24887.70 | 39.941 | *** | --- | --- |
|  | All taxa not in ratio 2 or 3 | 0.0770 | *A. nidulans*, *A. rugulosus* | 0.5994 | *A. unguis* | 0.0466 | -24864.04 | 47.440 | *** | 7.499 | ** |
|  | All taxa not in ratio 2 or 3 | 0.0784 | *A. nidulans*, *A. rugulosus* | 0.5967 | *A. unguis*, *A. versicolor* | 0.0545 | -24863.13 | 49.272 | *** | 9.331 | ** |
|  | All taxa not in ratio 2 or 3 | 0.0794 | *A. nidulans*, *A. rugulosus* | 0.5992 | *A. unguis*, *A. versicolor*,<br>*A. crustosus* | 0.0587 | -24863.11 | 49.312 | *** | 9.371 | ** |
|  | All taxa not in ratio 2 or 3 | 0.0801 | *A. nidulans*, *A. rugulosus* | 0.5975 | *A. unguis*, *A. versicolor*,<br>*A. crustosus*, *A. ustus* | 0.0631 | -24863.93 | 47.667 | *** | 7.726 | ** |
|  | All taxa not in ratio 2 or 3 | 0.0798 | *A. nidulans*, *A. rugulosus* | 0.5937 | *A. unguis*, *A. versicolor*,<br>*A. crustosus*, *A. ustus*,<br>*A. funiculosus* | 0.0673 | -24865.47 | 44.579 | *** | 4.638 | ** |
|  | All taxa not in ratio 2 or 3 | 0.0787 | *A. nidulans*, *A. rugulosus* | 0.5909 | *A. unguis*, *A. versicolor*,<br>*A. crustosus*, *A. ustus*,<br>*A. funiculosus*,<br>*A. conjunctus* | 0.0701 | -24866.67 | 42.186 | *** | 2.245 |  |
|  | All taxa not in ratio 2 or 3 | 0.0771 | *A. nidulans*, *A. rugulosus* | 0.5890 | *A. unguis*, *A. versicolor*,<br>*A. crustosus*, *A. ustus*,<br>*A. funiculosus*,<br>*A. conjunctus*,<br>*A. clavatoflavus* | 0.0723 | -24867.46 | 40.599 | *** | 0.658 |  |
|  | All taxa not in ratio 2 or 3 | 0.0730 | *A. nidulans*, *A. rugulosus* | 0.5876 | *A. clavatoflavus* | 0.1402 | -24865.09 | 45.344 | *** | 5.403 | ** |

| Taxon Set | Ratio 1 | $K_a/K_s$, 1 | Ratio 2 | $K_a/K_s$, 2 | Ratio 3 | $K_a/K_s$, 3 | LnL | D vs. 1-ratio | Sig.? | D vs. 2-ratio | Sig.? |
| --- | --- | --- | --- | --- | --- | --- | --- | --- | --- | --- | --- |
| C | All | 0.0718 | --- | --- | --- | --- | -20686.66 | --- | --- | --- | --- |
|  | All Pch2-retaining taxa | 0.0760 | All Pch2-lacking taxa | 0.0678 | --- | --- | -20685.70 | 1.927 |  | --- | --- |
|  | All Pch2-retaining taxa | 0.0760 | All Pch2-lacking taxa not in ratio 3 | 0.0722 | *A. unguis* | 0.0459 | -20683.14 | 7.055 | * | 0.358 |  |
|  | All Pch2-retaining taxa | 0.0760 | All Pch2-lacking taxa not in ratio 3 | 0.0752 | *A. unguis*, *A. versicolor* | 0.0536 | -20682.63 | 8.067 | * | 0.013 |  |
|  | All Pch2-retaining taxa | 0.0760 | All Pch2-lacking taxa not in ratio 3 | 0.0778 | *A. unguis*, *A. versicolor*,<br>*A. crustosus* | 0.0579 | -20682.89 | 7.549 | * | 0.055 |  |
|  | All Pch2-retaining taxa | 0.0759 | All Pch2-lacking taxa not in ratio 3 | 0.0618 | *A. unguis*, *A. versicolor*,<br>*A. crustosus*, *A. ustus* | 0.0824 | -20683.39 | 6.553 | * | 0.455 |  |
|  | All Pch2-retaining taxa | 0.0761 | All Pch2-lacking taxa not in ratio 3 | 0.0871 | *A. unguis*, *A. versicolor*,<br>*A. crustosus*, *A. ustus*,<br>*A. funiculosus* | 0.0654 | -20684.41 | 4.513 |  | 0.601 |  |
| D | All | 0.0699 | --- | --- | --- | --- | -20249.58 | --- | --- | --- | --- |
|  | All taxa not in ratio 2 | 0.0722 | *A. unguis* | 0.0457 | --- | --- | -20246.67 | 5.827 | * | --- | --- |
|  | All taxa not in ratio 2 | 0.0712 | *A. versicolor* | 0.0549 | --- | --- | -20248.66 | 1.840 |  | --- | --- |
|  | All taxa not in ratio 2 | 0.0704 | *A. crustosus* | 0.0629 | --- | --- | -20249.40 | 0.367 |  | --- | --- |
|  | All taxa not in ratio 2 | 0.0696 | *A. ustus* | 0.0726 | --- | --- | -20249.55 | 0.060 |  | --- | --- |
|  | All taxa not in ratio 2 | 0.0701 | *A. funiculosus* | 0.0670 | --- | --- | -20249.56 | 0.047 |  | --- | --- |
|  | All taxa not in ratio 2 | 0.0690 | *A. conjunctus* | 0.0854 | --- | --- | -20248.89 | 1.387 |  | --- | --- |
|  | All taxa not in ratio 2 | 0.0677 | *A. flavus* | 0.1060 | --- | --- | -20246.87 | 5.430 | * | --- | --- |
|  | All taxa not in ratio 2 | 0.0702 | *A. niger* | 0.0661 | --- | --- | -20249.52 | 0.121 |  | --- | --- |
|  | All taxa not in ratio 2 | 0.0705 | *A. terreus* | 0.0617 | --- | --- | -20249.30 | 0.561 |  | --- | --- |
|  | All taxa not in ratio 2 | 0.0702 | *A. clavatus* | 0.0651 | --- | --- | -20249.51 | 0.136 |  | --- | --- |
|  | All taxa not in ratio 2 | 0.0692 | *A. fumigatus* | 0.0804 | --- | --- | -20249.31 | 0.541 |  | --- | --- |
|  | All taxa not in ratio 2 | 0.0697 | *A. penicillioides* | 0.0722 | --- | --- | -20249.56 | 0.042 |  | --- | --- |
|  | All taxa not in ratio 2 | 0.0735 | *A. unguis, A. versicolor* | 0.0538 | --- | --- | -20246.35 | 6.468 | * | --- | --- |
|  | All taxa not in ratio 2 | 0.0741 | *A. unguis, A. versicolor, A. crustosus* | 0.0581 | --- | --- | -20246.78 | 5.593 | * | --- | --- |
|  | All taxa not in ratio 2 | 0.0745 | *A. unguis, A. versicolor, A. crustosus, A. ustus* | 0.0619 | --- | --- | -20247.52 | 4.115 | * | --- | --- |
|  | All taxa not in ratio 2 | 0.0740 | *A. unguis, A. versicolor, A. crustosus, A. ustus, A. funiculosus* | 0.0652 | --- | --- | -20248.49 | 2.176 |  | --- | --- |
|  | All taxa not in ratio 2 | 0.0721 | *A. unguis, A. versicolor, A. crustosus, A. ustus, A. funiculosus, A. conjunctus* | 0.0678 | --- | --- | -20249.32 | 0.531 |  | --- | --- |

Taxon Set A: *A. nidulans*, *A. rugulosus*, *A. unguis*, *A. versicolor*, *A. crustosus*, *A. ustus*, *A. funiculosus*, *A. conjunctus*, *A. flavus*, *A. niger*, *A. terreus*, *A. clavatus*, *A. fumigatus*, *N. fischeri*, *A. penicillioides*, *A. clavatoflavus*, *U. reesii* (outgroup)
Taxon Set B: set (A) without *A. nidulans* and *A. rugulosus*
Taxon Set C: *A. unguis*, *A. versicolor*, *A. crustosus*, *A. ustus*, *A. funiculosus*, *A. conjunctus*, *A. flavus*, *A. niger*, *A. terreus*, *A. clavatus*, *A. fumigatus*, *N. fischeri*, *A. penicillioides* (outgroup)
Taxon Set D: set (C) without *N. fischeri*
Note: Taxa in bold have no Pch2, underlined taxa have pseudogenized Msh4 and Msh5
Significance: blank = $p > 0.1$ , \* = $0.01 < p < 0.05$ , \*\* = $0.001 < p < 0.01$ , \*\*\* = $p < 0.001$ , --- = analysis not performed

**Table 4:** Codeml Analysis of Zip3 With Respect to Pch2 Loss.

| Ratio 1 | K<sub>a</sub>/K<sub>s</sub> 1 | Ratio 2 | K<sub>a</sub>/K<sub>s</sub> 2 | Ratio 3 | K<sub>a</sub>/K<sub>s</sub> 3 | LnL | D vs. 1-ratio | Sig.? | D vs. 2-ratio | Sig.? |
| --- | --- | --- | --- | --- | --- | --- | --- | --- | --- | --- |
| All | 0.0741 | --- | --- | --- | --- | -4688.99 | --- | --- | --- | --- |
| All intact Zip3 | 0.0531 | *A. nidulans*, *A. rugulosus* | 0.5802 | --- | --- | -4643.79 | 90.418 | *** | --- | --- |
| All intact Zip3 not in ratio 3 | 0.051 | *A. nidulans*, *A. rugulosus* | 0.5659 | *A. unguis* | 0.0734 | -4643.18 | 91.637 | *** | 1.219 |  |
| All intact Zip3 not in ratio 3 | 0.0528 | *A. nidulans*, *A. rugulosus* | 0.5787 | *A. versicolor* | 0.0587 | -4643.76 | 90.459 | *** | 0.041 |  |
| All intact Zip3 not in ratio 3 | 0.0531 | *A. nidulans*, *A. rugulosus* | 0.5803 | *A. crustosus* | 0.0536 | -4643.79 | 90.418 | *** | 0.001 |  |
| All intact Zip3 not in ratio 3 | 0.0554 | *A. nidulans*, *A. rugulosus* | 0.5745 | *A. conjunctus* | 0.0339 | -4643.07 | 91.840 | *** | 1.422 |  |
| All intact Zip3 not in ratio 3 | 0.052 | *A. nidulans*, *A. rugulosus* | 0.5804 | *A. flavus* | 0.068 | -4643.56 | 90.871 | *** | 0.453 |  |
| All intact Zip3 not in ratio 3 | 0.056 | *A. nidulans*, *A. rugulosus* | 0.5802 | *A. niger* | 0.0289 | -4642.62 | 92.743 | *** | 2.325 |  |
| All intact Zip3 not in ratio 3 | 0.05 | *A. nidulans*, *A. rugulosus* | 0.5828 | *A. terreus* | 0.0934 | -4642.54 | 92.905 | *** | 2.487 |  |
| All intact Zip3 not in ratio 3 | 0.054 | *A. nidulans*, *A. rugulosus* | 0.5799 | *A. clavatus* | 0.0382 | -4643.70 | 90.596 | *** | 0.178 |  |
| All intact Zip3 not in ratio 3 | 0.0521 | *A. nidulans*, *A. rugulosus* | 0.5813 | *A. fumigatus* | 0.0638 | -4643.64 | 90.713 | *** | 0.295 |  |
| All intact Zip3 not in ratio 3 | 0.0552 | *A. nidulans*, *A. rugulosus* | 0.5781 | *P. chrysogenum* | 0.0255 | -4643.06 | 91.860 | *** | 1.442 |  |
| All intact Zip3 not in ratio 3 | 0.0491 | *A. nidulans*, *A. rugulosus* | 0.5649 | *A. unguis*, *A. versicolor* | 0.0718 | -4642.76 | 92.460 | *** | 2.042 |  |
| All intact Zip3 not in ratio 3 | 0.049 | *A. nidulans*, *A. rugulosus* | 0.5723 | *A. unguis*, *A. versicolor*,<br>*A. crustosus* | 0.0629 | -4643.27 | 91.444 | *** | 1.026 |  |
| All intact Zip3 not in ratio 3 | 0.0563 | *A. nidulans*, *A. rugulosus* | 0.5821 | *A. unguis*, *A. versicolor*,<br>*A. crustosus*,<br>*A. conjunctus* | 0.0503 | -4643.65 | 90.689 | *** | 0.271 |  |

Taxon Set: *A. nidulans*, *A. rugulosus*, *A. unguis*, *A. versicolor*, *A. crustosus*, *A. conjunctus*, *A. flavus*, *A. niger*, *A. terreus*, *A. clavatus*, *A. fumigatus*, *P. chrysogenum* (outgroup)
Note: Taxa in bold have no Pch2, underlined taxa have pseudogenized Zip3
Significance: blank = $p > 0.1$ , \*\*\* = $p < 0.001$ , --- = analysis not performed

### Comparison of Zip3 K_a_/K_s_ Relative to Additional Cases of Msh4 and Msh5 Pseudogenization in Additional Eurotiomycetes

While performing the above Codeml analyses, bioinformatic searches for the four meiosis genes of interest were performed in the genomes of four additional Eurotiomycetes (*Eurotium herbariorum, Penicillium chrysogenum, Penicillium marneffei, Talaromyces stipitatus*; see supplemental file S1 for genome information) with the intent to potentially utilize orthologs from an outgroup more closely related to genus *Aspergillus* than *U. reesii*. Unexpectedly, though, all four of these species were found to have undergone independent losses of at least one of the four genes of interest: Pch2 was found in only *E. herbariorum,* while Msh4 and Msh5 were each pseudogenized in *E. herbariorum* and *T. stipitatus* (Figure 1, Figure 4, Figure 5, Figure 7, Table 1). Zip3 was not unambiguously pseudogenized in any of these four species; however, three of the four sequences had some notable characteristics. *P. marneffei* and *T. stipitatus* share two in-frame deletions (Figure 10A) that generate a somewhat shorter predicted Zip3p sequence than a representative full-length intact Zip3 ortholog (Table 1). *T. stipitatus* Zip3 has two additional in-frame deletions within the fourth exon (not adjacent to intron boundaries) in a moderately conserved region of the gene (Figure 10B). *E. herbariorum* Zip3 has at least one possible loss-of function mutation: a putative GT⃗AT substitution predicted to disrupt the 5’ splice site of the first intron (highly conserved with other manually annotated Eurotiomycete Zip3 orthologs.) Two additional possible ORF disruptions exist near the fourth intron; if one hypothesized a single frameshift mutation, the downstream sequence in the shifted frame more closely matches that of intact sequences and the fourth intron position is better conserved—albeit with a disruption to the 3’ splice site of that intron (GT..AC, Figure 11). Because the possibility cannot be excluded that an alternative upstream GC..AG splice site is utilized that maintains an open reading frame, the phylogenetic analyses utilized the more conservative annotation with a single ORF disruption (the first intron GT⃗AT, Figure 11). Either annotation is consistent with *E. herbariorum* Zip3 having very recently pseudogenized; however, genome sequence trace reads were not available for examination to exclude the possibility of local poor coverage or sequencing errors.

**Figure 10:**
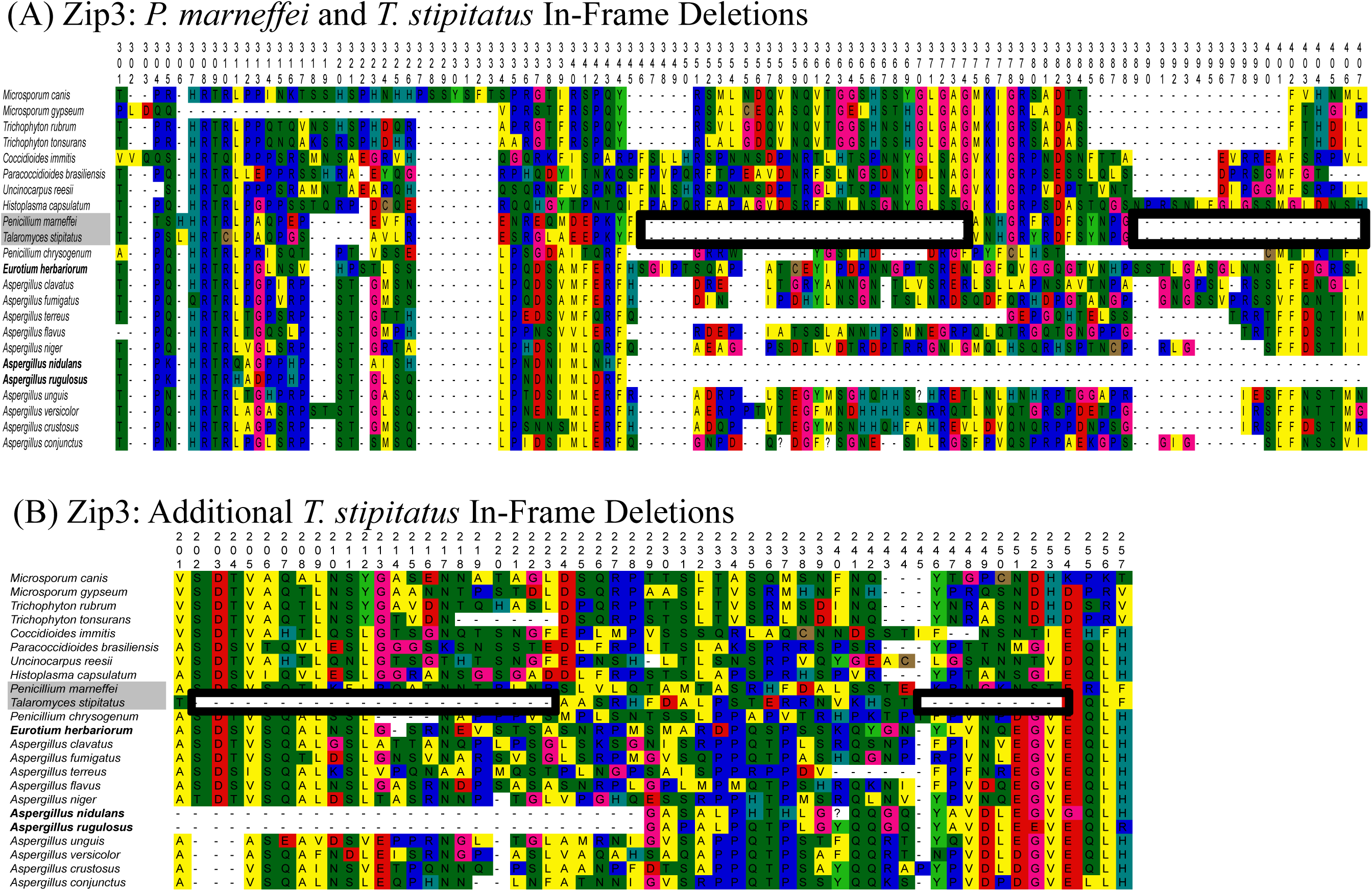
*T. stipitatus* and *P. marneffei* Zip3 In-Frame Deletions. This excerpt of the Zip3 amino acid multiple sequence alignment shows (A) homologous in-frame deletions in *T. stipitatus* and *P. marneffei* and (B) additional in-frame deletions specific to *T. stipitatus*. Numbers indicate the position in the full alignment (will be in supplemental file S4; available on request in the interim on BioRxiv.)

**Figure 11:**
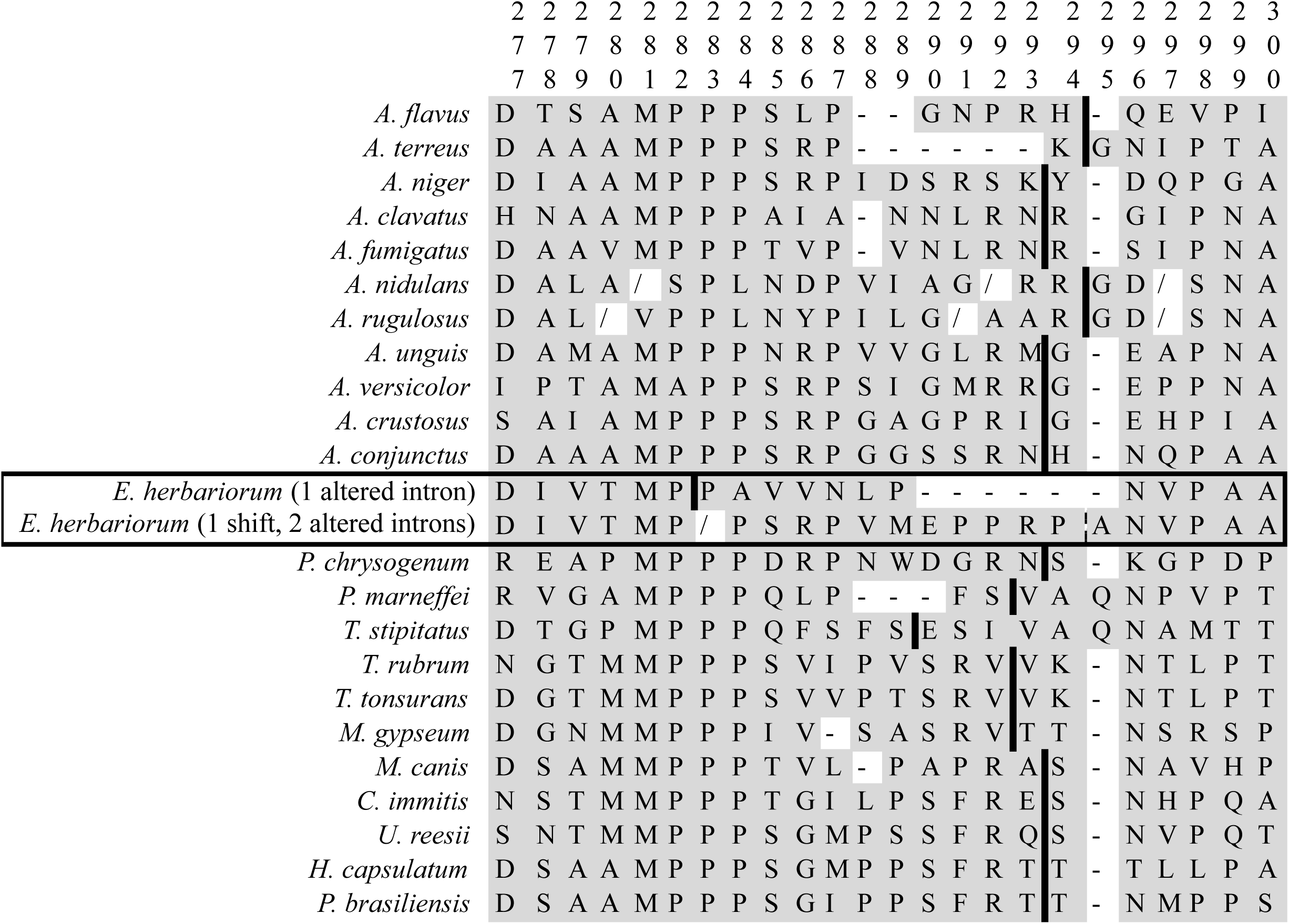
*E. herbariorum* Zip3 Annotation Comparison. This excerpt of the Zip3 amino acid multiple sequence alignment compares the more conservative *E. herbariorum* annotation used in phylogenetic analysis (“1 altered intron”) with an alternative annotation that would suggest more disruptions to *E. herbariorum* Zip3 (“1 shift, 2 altered introns”). Thick black lines represent a phase 1 intron, and the broken black and white vertical line in the second *E. herbariorum* sequence is a putative former phase 1 intron with a splice site alteration. Frameshifts are noted by “/” marks. Numbers indicate the position in the full alignment (will be in supplemental file S4; available on request in the interim on BioRxiv.)

To determine whether the atypical sequence features of *E. herbariorum, P. marneffei*, and/or *T. stipitatus* Zip3 orthologs were associated with gene-wide K_a_/K_s_ elevation (potentially indicative of on-going or incipient pseudogenization), additional Codeml trials were performed (Table 5). Weakly significant support was seen for a model in which *E. herbariorum* Zip3 has a separate, higher K_a_/K_s_ (0.2013 *vs.* 0.0490, 0.01<p<0.05, Table 5) compared to the clearly intact Eurotiomycete Zip3 orthologs (all of which except *U. reesii* were found to not have K_a_/K_s_ values substantially different from each other in Table 4). This is again consistent with very recent pseudogenization of *E. herbariorum* Zip3 and a lack of time for the K_a_/K_s_ ratio to approach the expected eventual value of 1. The tests examining *T. stipitatus* and *P. marneffei* Zip3 (added singly or together to the core “intact Zip3” taxon set in Table 5) were more ambiguous because some K_s_ estimates for the additional test species were markedly higher or lower than expected (*i.e.,* 57.5289, 3.5610, 0; Table 5). This problem persisted regardless of whether the tree was rooted or unrooted and may be due to the necessary placement of *T. stipitatus* and *P. marneffei* (sister taxa) as the most basal branch other than the distantly related—but closest available— exemplar Onygenales outgroup species *U. reesii* (van den Berg et al., 2008; Wang et al., 2009). The two cases in which K_s_ estimates were not atypically high or low (taxon set D, *P. marneffei* assigned separate ratio; taxon set D, *P. marneffei*, *T. stipitatus*, and uniting branch assigned separate ratio) supported an elevated Zip3 K_a_/K_s_ in the species or clade assigned a separate ratio (Table 5). These results are suggestive of changes in sequence evolution constraints in *P. marneffei* and *T. stipitatus* Zip3, although the functional implications (if any) are unclear.

**Table 5:** Codeml Analysis of Zip3 With Respect to Msh4 and Msh5 Pseudogenization.

| Taxon Set | Ratio 1 | K<sub>a</sub>/K<sub>s</sub> 1 | Ratio 2 | K<sub>a</sub>/K<sub>s</sub> 2 | LnL | D vs. 1-ratio | Sig.? | Ratio 2 K<sub>a</sub>, K<sub>s</sub> |
| --- | --- | --- | --- | --- | --- | --- | --- | --- |
| A | All | 0.0512 | --- | --- | -4602.49 | --- | --- | --- |
| A | All not in ratio 2 | 0.0490 | *E. herbariorum* | 0.2013 | -4599.82 | 5.349 | * | 0.0786, 0.3905 |
| B | All | 0.0467 | --- | --- | -4705.36 | --- | --- | --- |
| B | All not in ratio 2 | 0.0486 | *P. marneffeii* | 0.0026 | -4702.87 | 4.983 | * | 0.1500, 57.5289 |
| C | All | 0.0476 | --- | --- | -4695.95 | --- | --- | --- |
| C | All not in ratio 2 | 0.0479 | *T. stipitatus* | 0.0404 | -4695.92 | 0.054 |  | 0.1440, 3.5610 |
| D | All | 0.0540 | --- | --- | -4992.89 | --- | --- | --- |
| D | All not in ratio 2 | 0.0487 | *P. marneffeii* | 0.331 | -4986.79 | 12.193 | ** | 0.0509, 0.1537 |
| D | All not in ratio 2 | 0.0493 | *T. stipitatus* | 999 | -4985.98 | 13.815 | ** | 0.0379, 0 |
| D | All not in ratio 2 | 0.0471 | *P. marneffeii*,<br>*T. stipitatus* | 0.1154 | -4987.70 | 10.379 | * | *T. stipitatus*: 0.0293, 0.2536;<br>*P. marneffeii*: 0.0441, 0.3821;<br>Uniting branch: 0.1231, 1.0670 |

Core Taxa: *A. unguis*, *A. versicolor*, *A. crustosus*, *A. conjunctus*, *A. flavus*, *A. niger*, *A. terreus*, *A. clavatus*, *A. fumigatus*, *P. chrysogenum*, *U. reesii* (outgroup)
Taxon Set A: Core plus *E. herbariorum*
Taxon Set B: Core plus *P. marneffeii*
Taxon Set C: Core plus *T. stipitatus*
Taxon Set D: Core plus *P. marneffeii* and *T. stipitatus*
Significance: blank = $p > 0.1$ , \* = $0.01 < p < 0.05$ , \*\* = $0.001 < p < 0.01$ , --- = analysis not performed

### Comparison of Known or Predicted Homothallism to ZMM Pseudogenization Distribution

Literature reports that two of the species with pseudogenized Msh4 and Msh5 were also homothallic (*A. nidulans* (Galagan et al., 2005), *E. herbariorum* (Dyer and O’Gorman, 2012)) and, conversely, several species with intact Msh4 and Msh5 were heterothallic (*A. flavus* (Horn et al., 2009a), *A. terreus* (Arabatzis and Velegraki, 2013), *A. fumigatus* (O’Gorman et al., 2009)) prompted a BLAST survey of MAT1-1-1 and MAT1-2-1 distribution in the species for which genome sequences were publicly available. Interestingly, all three species with pseudogenized Msh4 and Msh5 that had searchable genome sequences had both MAT1-1-1/MAT1 and MAT1-2-1/MAT2 (implying homothallism (Dyer and O’Gorman, 2011)), while most of the closest available relatives retaining Msh4 and Msh5 (*i.e.*, *A. versicolor* and *A. sydowii* relative to *A. nidulans*, *P. marneffei* relative to *T. stipitatus*) had only MAT1-1-1 or only MAT1-2-1 (implying heterothallism) (Figure 12). An exception was that both *E. herbariorum* and *A. glaucus* (sister taxa relative to other examined Eurotiomycetes (Peterson, 2008; Hubka *et al*., 2013)) had MAT1-1-1 and MAT1-2-1 orthologs detected (Figure 12) despite *A. glaucus* having apparently intact orthologs of Msh4, Msh5, and Zip3 (Figure 1). Another known homothallic species, *Neosartorya fischeri* (Rydholm *et al*., 2007), also had intact orthologs of all four sought meiosis genes (Figure 1). These results indicate that, while all of the Eurotiomycetes with pseudogenized ZMM genes are (or are likely to be) homothallic, not all homothallic Eurotiomycetes have pseudogenized ZMM genes.

**Figure 12:**
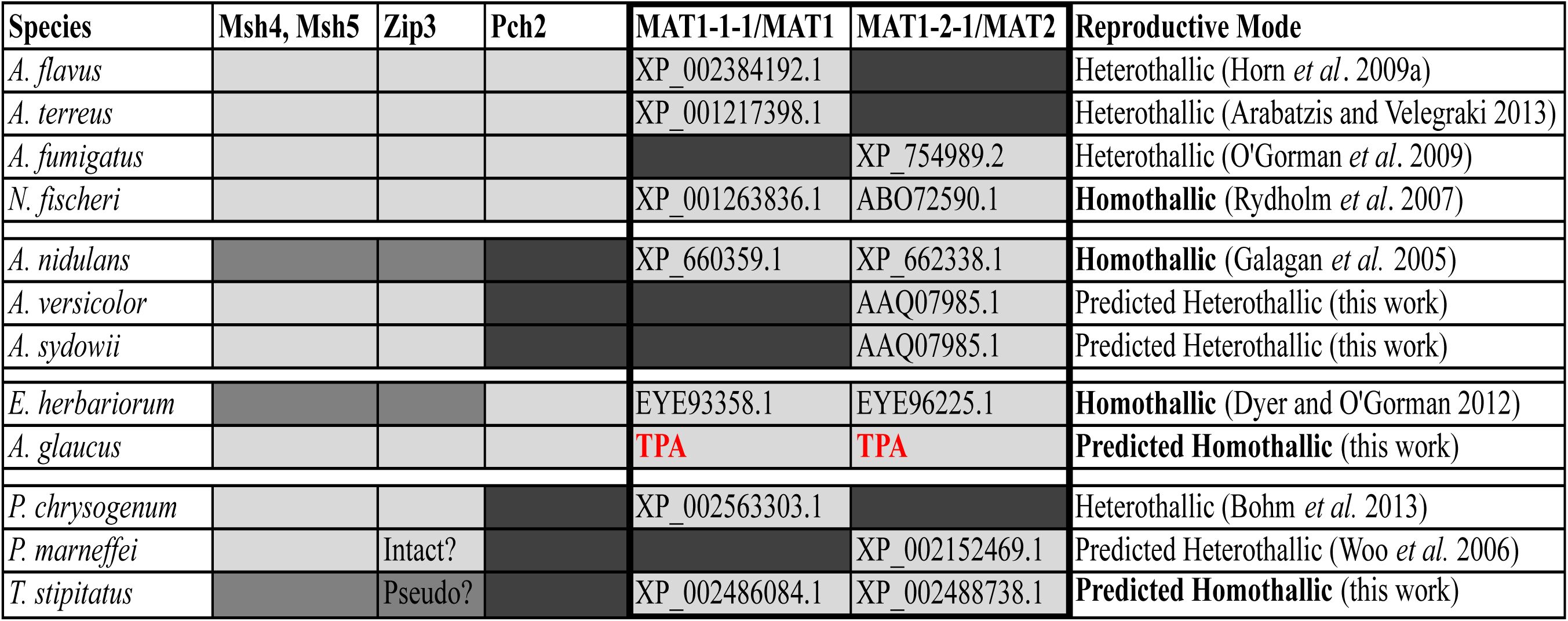
Msh4, Msh5, and Zip3 Losses in Species with MAT1 and MAT2. Light gray boxes are intact orthologs, medium gray boxes are pseudogenized orthologs, and dark gray boxes indicate that an ortholog was not detected. Bold font for MAT accession numbers indicates new predicted protein sequences from this work. A reproductive mode of “predicted homothallic” or “predicted heterothallic” indicates that sexual reproduction has not been reported yet in that species to our knowledge at the time of writing.

## Discussion

### Comparisons to Previous Studies and Methodological Comments

The substantial degradation of the *A. nidulans* Msh4 and Msh5 pseudogenes (Table 1) explains the previous inability of Woo *et al*. (2006) to identify these genes in *A. nidulans*, although that study reported only retrieving protein sequences already in the NCBI databases at the time and not TBLASTN or BLASTP examining the *A. nidulans* genome itself. Because inability to detect an ortholog by bioinformatics methods can occur for many reasons other than gene loss (*e.g.*, genome sequence coverage gaps, assembly errors, rapid sequence evolution) these degraded pseudogenes are rare cases where detection failure can be directly confirmed as gene loss. The present study is also consistent with NCBI searches by Wu and Burgess (2006) that did not identify an *A. nidulans* Pch2 ortholog using *C. elegans* and *S. cerevisiae* BLASTP queries. The distribution of Zip3 homologs in *Mus musculus* (Ward *et al*., 2007; Reynolds *et al*., 2013), *Arabidopsis thaliana* (Chelysheva et al., 2012), and *S. cerevisiae* (Agarwal and Roeder, 2000) predicted that Zip3 would have been ancestrally present in the MRCA of fungi as a whole. However, Zip3 orthologs had not been previously sought in Eurotiomycetes save for two exceptions: Desjardins *et al*. (2011) sought but did not identify Zip3 orthologs in four Eurotiomycete species, while Chelysheva *et al*. (2012) reported a putative *P. marneffei* homolog. The present work predicts a relatively broad distribution of Zip3 among fungi that may parallel the distribution of Msh4 and Msh5.

In each case, considering flanking gene synteny conservation bypassed two major limitations that impact the study of genes with precedent for derived losses. First, attempting degenerate PCR directly on the gene of interest would have carried a high risk of failure for orthologs that were rapidly evolving, pseudogenizing, or lost. Second, bioinformatics searches using orthologs of the target genes of interest as queries are not able to easily identify short, highly degraded pseudogene remnants depending on the degree of degradation; the synteny context provides corroborating evidence that a pseudogene remnant has been accurately identified as opposed to a spurious, low E-value alignment with a non-orthologous sequence. The reliance on synteny did produce some limitations in the degenerate PCR work, though.

First, the failure to amplify the predicted Zip3-containing region from four species could plausibly be explained by flanking gene synteny changes—changes that were relatively common on the 3’ flanking side of Zip3 in exemplar Eurotiomycete genome sequences (Figure 2). Second, synteny changes surrounding *A. penicillioides* Pch2 (whether by inversion, translocation, or additional gene insertions) could explain the inability to amplify sequence between the Rcy1 fragment and the Pch2-Cyt1 region. Serendipitous primer binding to the 5’ end of Pch2 allowed amplification of the *A. penicillioides* ortholog; however, had this serendipitous binding not occurred, the lack of amplification would have given an ambiguous result that would not have allowed inferences about Pch2 status. Third, an apparent second copy of Hsp78 in *A. funiculosus* was amplified through off-target primer binding. Although this did not preclude the ability to amplify the *A. funiculosus* Hsp78-Msh5-Sas10 region, it illustrates how this approach would be more challenging to apply to target genes in which at least one flanking gene has multiple copies.

### Number of and Support for Losses

#### Msh4 and Msh5

The *A. nidulans* and *A. rugulosus* orthologs of Msh4 and Msh5 contain in-frame stop codons, frameshift mutations, loss of start codons, loss of consensus splice site sequences, and/or large deletions (Table 1)—some of which are present at homologous positions (Figure 3); this provides molecular evidence consistent with loss of functional Msh4p and Msh5p in the MRCA of these two species. The wild type meiosis of *A. nidulans* lacks crossover interference (Strickland, 1958; Egel-Mitani *et al*., 1982) and, by extension, class I crossovers (Lynn *et al*., 2007); this suggests that the loss of class I crossover-related genes in *A. nidulans* was not associated with exaptation of existing genes or acquisition of novel genes that provide the same class I crossover formation function (*i.e.*, loss of these ZMM genes was also associated with a loss of a feature in meiosis.) A parallel situation exists in *S. pombe*, another fungus that has independently lost Msh4 and Msh5 (Malik *et al*., 2008) along with class I crossover formation and crossover interference (Snow, 1979; Hollingsworth and Brill, 2004). These prior studies, coupled with the consistent defects in class I crossover formation from Msh4 and/or Msh5 knockout mutants in other model organisms (Kneitz *et al*., 2000; Novak *et al*., 2001; de los Santos *et al*., 2003; Argueso *et al*., 2004; Higgins *et al*., 2004; Lynn *et al*., 2007; Higgins *et al*., 2008), generate the prediction that *E. herbariorum* and *T. stipitatus* also lack class I crossovers and crossover interference. These latter two species with pseudogenized Msh4 and Msh5 were consistently supported as each having a sister taxon with intact orthologs of these genes (*T. stipitatus* with *P. marneffei*, *E. herbariorum* with *A. penicillioides*; Figures 4-6), indicating that Msh4 and Msh5 have independently pseudogenized in three Eurotiomycete lineages.

#### Zip3

Zip3 was clearly pseudogenized in *A. nidulans* and *A. rugulosus* (Table 1), with homologous candidate loss-of-function mutations (Figure 3) again implying its pseudogenization in the MRCA of these two species. Early pseudogenization of *E. herbariorum* Zip3 seems likely given its elevated K_a_/K_s_ relative to clearly intact orthologs (Table 5) and the presence of at least one ORF disruption. However, the possibility of alternative splicing around that region resulting in a protein that is functional (albeit with some relaxation of sequence evolution constraint) cannot be excluded without further exploration of *E. herbariorum* Zip3 transcripts during meiosis. The functional status of the *T. stipitatus* and *P. marneffei* Zip3 orthologs is difficult to predict because all deletions in these orthologs were in-frame (Figure 10) and most of the K_a_/K_s_ analyses had problematic estimates of K_s_ (Table 5). The presence of additional in-frame deletions in the Zip3 of *T. stipitatus* relative to *P. marneffei* (Figure 10) could be consistent with incipient pseudogenization of Zip3 associated with pseudogenization of Msh4 and Msh5, as seen in the first two cases (*A. nidulans-A. rugulosus*, *E. herbariorum*). However, an alternative explanation could be positive selection on some regions of *T. stipitatus* Zip3 associated with compensatory changes secondary to the Msh4 and Msh5 losses. A third possibility emerges if Eurotiomycete Zip3 orthologs were to share the same synaptonemal complex assembly functions as the *S. cerevisiae* ortholog (Agarwal and Roeder, 2000; Serrentino *et al*., 2013); in this scenario, loss of Msh4 and Msh5 in *T. stipitatus* could relax selection on any Zip3p domains crucial for its established interaction with Msh5p (Agarwal and Roeder, 2000) while leaving domains relevant to any Msh4p- and Msh5p-independent functions under purifying selection. However, the specific regions of Zip3p most crucial for Msh5p interaction or synapsis functions in *S. cerevisiae* have not been characterized to our knowledge, and the functions of specific sequence regions of Zip3 in Eurotiomycetes are also yet unknown.

The phylogenetic distribution of synaptonemal complex (SC)-related functions in Zip3 is rather disparate, and these functions are variable in nature. No synapsis-related functions have been reported for plant or mouse Hei10 (Ward *et al*., 2007; Chelysheva *et al*., 2012; Wang *et al*., 2012); by contrast, *S. cerevisiae* Zip3p functions in SC assembly (Agarwal and Roeder, 2000), *C. elegans* ZHP-3 is needed for SC disassembly but not SC assembly (Bhalla *et al*., 2008), and *M. musculus* RNF212 functions in X/Y synapsis (Reynolds *et al*., 2013). The SC of Eurotiomycetes has not been investigated in species other than *A. nidulans* to our knowledge, but *A. nidulans* is reported to lack a canonical SC (Egel-Mitani et al., 1982). Whether the loss of canonical SC in *A. nidulans* is associated directly with loss of Zip3 (or, indeed, whether other Eurotiomycetes exhibit canonical SC formation) remains to be determined. An indirect hint potentially could be in the observation that the Msh4 and Msh5 pseudogenes in *A. nidulans, A. rugulosus, E. herbariorum,* and *T. stipitatus* are all more degraded (more premature stop codons and frameshifts, more numerous or larger deletions, *etc.*) than the Zip3 orthologs in these species (Table 1). One possible explanation for this pattern could be if Zip3 is retained (or, in *A. nidulans* and *A. rugulosus*, was initially retained before later pseudogenization) in association with some unknown Msh4- and Msh5-independent functions, whether the SC or otherwise; however, an alternative explanation is that chance differences in the size of deletion mutations are sufficient to explain the difference in amounts of recognizable sequence in the *A. nidulans* and *A. rugulosus* Zip3 pseudogenes relative to the Msh4 and Msh5 pseudogenes. Regardless of this speculation about possible SC functions of Eurotiomycete Zip3, all functionally characterized Zip3 homologs exhibit roles in class I crossover formation (Agarwal and Roeder, 2000; Borner *et al*., 2004; Ward *et al*., 2007; Bhalla *et al*., 2008; Chelysheva *et al*., 2012; Wang *et al*., 2012; Reynolds *et al*., 2013) and many are known to have close functional and physical relationships with Msh4 and/or Msh5 (Agarwal and Roeder, 2000; Borner *et al*., 2004; Chelysheva *et al*., 2012; Reynolds *et al*., 2013). Zip3 also has been found to show substantial sequence coevolution with Msh4, Msh5, and other class I crossover-related genes in Saccharomycotina fungi (Clark *et al*., 2013). Collectively, these prior findings support interpretation of the present Eurotiomycete results as further evidence of the functional association (and molecular coevolution) of Zip3 with Msh4 and Msh5, in which Zip3 was pseudogenized or showing other signs of possible relaxed selection in all four species with pseudogenized Msh4 and Msh5.

#### Pch2

In contrast to the Msh4, Msh5, and Zip3 losses (in which pseudogenes are still detectable), all of the putative Pch2 losses are based on an inability to identify an ortholog in publicly available genome sequences and/or at the expected syntenic location (Figure 1, Figure 2). Prolonged neutral evolution of a pseudogene is expected to eventually result in a sequence becoming unrecognizable as an ortholog due to accumulation of nonsynonymous changes and/or deletions. Although translocation of intact Pch2 without either of the consensus flanking genes (*i.e.*, present but not amplifiable by flanking gene-targeted PCR) cannot be formally excluded for the species examined only by degenerate PCR, there was not a precedent for this in the taxa with genome sequences assessed for both synteny and Pch2 ortholog queries—when Pch2 was found in these taxa, it was always single-copy and flanked by at least one of Rcy1 and Cyt1 (usually both; Figure 2.) Pch2 was also undetected in genome-wide TBLASTN searches of *A. nidulans*, *A. versicolor*, and *A. sydowii* (a clade including *A. rugulosus* and *A. unguis* [Figure 1]) as well as *A. zonatus* (sister taxon to *A. clavatoflavus*.) Similarly, while sequencing coverage gaps and/or assembly errors could potentially explain inability to detect Pch2 in a particular genome, invoking these explanations for all seven Eurotiomycete genome sequences apparently lacking Pch2 is not parsimonious (*i.e.*, coverage gaps and assembly errors would need to occur at the same location for the same gene in seven independently assembled genomes.) The TBLASTN parameters were also sufficient to return numerous genomic DNA hits more similar to nonorthologous AAA ATPases than to Pch2 in all examined Eurotiomycetes, regardless of whether a Eurotiomycete or *S. cerevisiae* query was used (data not shown). This suggests that the lack of detected Pch2 was probably not due to overly stringent E thresholds or substantial dissimilarity between the query and target species sequences. Therefore, the cases in which Pch2 orthologs were not detected are likely to represent derived gene losses. This present work’s TREE-PUZZLE analyses consistently rejected both tested topologies depicting *A. clavatoflavus* as a sister taxon to the *A. nidulans-A. conjunctus* clade (Figure 9) and therefore support two independent Pch2 losses in genus *Aspergillus*. The phylogenetic placements of *P. chrysogenum*, *P. marneffei*, and *T. stipitatus* relative to Pch2-retaining Eurotiomycetes in both this study’s gene trees (Figure 1, Figures 4-6) and other reports (van den Berg *et al*., 2008) suggest that two more independent Pch2 losses have occurred among these species: one in *P. chrysogenum* and one in the MRCA of *P. marneffei* and *T. stipitatus*. This total of four independent losses makes Pch2 the most frequently lost of the four examined meiosis genes in these Eurotiomycetes (three independent losses for Msh4 and Msh5, at least one for Zip3).

Unlike Msh4 and Msh5, where broadly conserved knockout phenotypes in model species ordinarily retaining the genes generate a clear phenotype prediction similar to wild type meiosis of species lacking the genes (*i.e.*, lack of class I crossovers in *S. cerevisiae msh4*Δ and *msh5*Δ mutants mirrors wild type *A. nidulans* and *S. pombe* (Strickland, 1958; Snow, 1979; Egel-Mitani *et al*., 1982; Argueso *et al*., 2004; Malik *et al*., 2008; Lynn *et al*., 2007; Hollingsworth and Brill, 2004)), predicting or identifying the effects of the ancestral Pch2 loss on wild type *A. nidulans* meiosis is challenging for several reasons. First, Pch2 functions have not been characterized in Eurotiomycetes and have been found to vary across the model species in which orthologs have been characterized. *C. elegans* Pch-2 functions in meiotic synapsis checkpoint activity (Bhalla and Dernburg, 2005), *D. melanogaster* PCH2 functions in checkpoint delays independent of synapsis (Joyce and McKim, 2009), while *M. musculus* TRIP13 is required for DSB repair itself and yet-unknown somatic functions (Li and Schimenti, 2007). *S. cerevisiae* Pch2 has been implicated in diverse facets of meiosis: crossover interference (Joshi *et al*., 2009), frequency of intersister *vs.* interhomolog recombination (Zanders *et al*., 2011), synaptonemal complex protein localization (San-Segundo and Roeder, 1999), and gene conversion frequency regulation (Zanders and Alani, 2009); two additional processes affected in backgrounds with specific additional mutations include pachytene arrest (San-Segundo and Roeder, 1999) and DNA double-strand break (DSB) formation (Farmer *et al*., 2012). Without knowing the ancestral Eurotiomycete function of Pch2, one cannot yet confidently predict which features of wild type *A. nidulans* meiosis may be due to the Pch2 loss. Second, one of the effects of *pch2*Δ in *S. cerevisiae* is a weakening of crossover interference (Joshi *et al*., 2009); even if one assumes this function to be conserved in Eurotiomycete Pch2, the effect of lacking Pch2 in *A. nidulans* could not be disentangled from the loss of Msh4 and Msh5 (which is expected to eliminate all crossovers that exhibit interference and thus should eliminate observations of crossover interference overall (Lynn *et al*., 2007)). Third, knockout phenotypes of Pch2 orthologs in most tested model species are typically mild, with deleterious phenotypes generally apparent only in backgrounds with loss of function in additional meiosis genes (San-Segundo and Roeder, 1999; Bhalla and Dernburg, 2005; Wu and Burgess, 2006; Joyce and McKim, 2009; Zanders and Alani, 2009; Zanders *et al*., 2011). Therefore, loss of Pch2 alone may not necessarily have conferred an initial obvious change to the dynamics of wild type meiosis. Notably, Zanders and Alani (2009) found that *S. cerevisiae pch2*Δ*msh5*Δ double mutants had lower spore viability (26%) than single mutants of *pch2*Δ (95%) or *msh5*Δ (36%); this predicts that prior loss of Pch2 would not have conferred an immediate amelioration of the effects of losing Msh5 (see next subsection for further discussion.) Functional characterization of meiosis genes in Eurotiomycetes with intact orthologs and inducible sexual reproduction will be necessary to more reliably identify ways in which the overall process of meiosis in Pch2-lacking Eurotiomycetes has been changed (if at all) and how Pch2 loss is so frequently tolerated despite its presence still being broadly conserved.

### Possible Factors Influencing Meiosis Gene Loss in Eurotiomycetes

The multiple independent meiosis gene losses in the examined Eurotiomycetes raise the question of what molecular, genetic, ecological, or environmental properties are most conducive to tolerating losses of these genes. Some exemplar factors that will be discussed are: loss of sexual reproduction, loss of Pch2, homothallism, variation in class I and class II crossover frequency, chromosome number and size, and temperature during sexual reproduction.

Loss of sexual reproduction and transitioning to exclusive asexuality is predicted to relax selection on genes that function exclusively in meiosis, leading to their pseudogenization and loss (Schurko & Logsdon, 2008). However, all of the Eurotiomycetes identified with ZMM pseudogenes in this work are reported to experience sexual reproduction (Butinar *et al*., 2005; Todd *et al*., 2007; Peterson, 2008; Lopez-Villavicencio *et al*., 2010). Sexual *Aspergillus* species are also known within clades with Pch2 (Horn *et al*., 2009a; O’Gorman *et al*., 2009) and without Pch2 (Fennell and Raper, 1955). This indicates that the loss of Msh4, Msh5, Zip3, or Pch2 does not coincide with loss of sexual reproduction in Eurotiomycetes.

The finding that Pch2 loss preceded loss of Msh4, Msh5, and Zip3 in the *A. nidulans-A. conjunctus* clade raised the question of whether prior loss of Pch2 could have contributed in some way to the subsequent loss events in that clade or in other Eurotiomycetes. However, the present results do not support prior Pch2 loss as a strong, direct influence on loss of Msh4, Msh5, or Zip3 in the examined Eurotiomycetes. First, no consistent significant difference in Msh4, Msh5, or Zip3 K_a_/K_s_ was found between Pch2-lacking and Pch2-retaining species in any of the many Codeml trials (Tables 2-4). Second, loss of Pch2 did not always lead to subsequent loss of the other three genes; Msh4 and Msh5 remained intact (Figure 1) and under purifying selection (Tables 2-3) in most of the Pch2-lacking Eurotiomycete species. Finally, *E. herbariorum* experienced pseudogenization of Msh4, Msh5, and (likely) Zip3 despite Pch2 being present and intact (Figure 1). The observation that two of the three identified Msh4 and Msh5 pseudogenization cases were preceded by Pch2 loss (*A. nidulans-A. rugulosus*, *T. stipitatus*) remains intriguing nonetheless. One possibility could be that loss of Pch2 in Eurotiomycetes confers changes in genetic background that can contribute to tolerance of subsequent meiosis gene losses (while being neither strictly necessary nor alone sufficient for additional gene losses.) However, this scenario would require differences from *S. cerevisiae*, where *pch2*Δ*msh5*Δ have lower spore viability than *msh5*Δ single mutants (Zanders and Alani, 2009) such that Pch2 loss further compounds the deleterious meiosis phenotype of Msh5 loss; further characterization of Eurotiomycete Pch2 would be necessary to evaluate similarities to or differences from *S. cerevisiae* Pch2. Regardless, prior loss of Pch2 does not seem to be immediately associated with substantial changes in the sequence evolution constraints of Msh4, Msh5, and Zip3.

The most striking commonality among the independent Msh4 and Msh5 pseudogenization events is that all involve taxa known to be homothallic (*A. nidulans* (Galagan et al., 2005), *E. herbariorum* (Dyer and O’Gorman, 2012)) or appear to be potentially homothallic based on the presence of both MAT1-1-1/MAT1 and MAT1-2-1/MAT2 orthologs within a single strain’s genome sequence (*T. stipitatus*, Figure 12). (At the time of writing, confirmation of *A. rugulosus* as homothallic or heterothallic has not been published to our knowledge.) By contrast, in two of the three independent Msh4 and Msh5 loss cases, the closest sister taxon with intact Msh4 and Msh5 and a publicly available genome sequence (*A. versicolor* and *A. sydowii* relative to *A. nidulans* and *A. rugulosus*, *P. marneffei* relative to *T. stipitatus*) had only MAT1-1-1 or MAT1-2-1 detected (Figure 12); this result implies likely heterothallism (obligate mating with individuals of two different mating types) in these sister species (Whittle *et al*., 2011). Msh4 and Msh5 pseudogenization in Eurotiomycetes therefore appears to be strongly associated with homothallism. Homothallism is the ability to reproduce with isolates of the same mating type, allowing for self-fertilization and inbreeding in addition to outcrossing or opposite mating type crosses (Whittle *et al*., 2011). Inbreeding reduces effective population size (N_e_), reduces the efficacy of selection, and increases the probability of alleles becoming fixed by genetic drift if they arise in a population or were present before the reduction in N_e_ (reviewed in Whittle *et al*., 2011). Under these conditions, a loss-of-function mutation in Msh4 or Msh5 could have a greater likelihood of becoming fixed (compared to conditions with a larger N_e_) if loss of class I crossovers was initially either neutral or only mildly deleterious; if the loss of class I crossovers was severely deleterious, as in *S. cerevisiae* (Novak et al., 2001), nonfunctional Msh4 or Msh5 alleles would not be expected to persist. Although homothallism by itself would not directly affect the severity of a loss-of-function mutation, the resulting reduced effective population size from inbreeding could both temporarily dampen the severity of deleterious alleles and accelerate their fixation relative to the chances of fixation with a larger effective population size. The specific molecular or environmental conditions that influence whether loss of class I crossovers would be highly deleterious or only mildly deleterious are yet unknown and are not necessarily uniform for all homothallic Eurotiomycetes. For example, *N. fischeri* is homothallic (Rydholm *et al*., 2007) and the *A. glaucus* genome sequence contains putative orthologs of both MAT1-1-1/MAT1 and MAT1-2-1/MAT2 (Figure 12), but both species have intact orthologs of all four sought meiosis genes (Figure 1). Another key point is that, unlike Msh4 and Msh5, the Pch2 trends do not closely mirror homothallism or heterothallism status in Figure 12. One possible explanation for this difference could be from Pch2 loss being nearly neutral in Eurotiomycetes, as observed in many model organisms (San-Segundo and Roeder, 1999; Bhalla and Dernburg, 2005; Wu and Burgess, 2006; Joyce and McKim, 2009; Zanders and Alani, 2009; Zanders *et al*., 2011); if this is also the case in Eurotiomycetes, Pch2 losses potentially could become fixed even without the lower N_e_ from homothallism (which would have a relatively greater influence on mutations that were initially mildly deleterious.) Overall, homothallism alone does not universally lead to meiosis gene losses. However, homothallism in combination with additional predisposing factors for tolerance of class I crossover loss (possible candidates discussed below) could be a plausible contributing mechanism for the observed Msh4 and Msh5 pseudogene distribution in Eurotiomycetes.

Loss of class I crossovers through loss-of-function mutations in Msh4 and/or Msh5 orthologs in model species such as *S. cerevisiae*, *C. elegans*, and *A. thaliana* means that (i) fewer crossovers form and (ii) any remaining crossovers are class II (Msh4- and Msh5-independent, Mus81- and Eme1/Mms4-dependent) and thus do not exhibit crossover interference (Zalevsky *et al*., 1999; Kelly *et al*., 2000; de los Santos *et al*., 2003; Higgins *et al*., 2004; Hollingsworth and Brill, 2004; Lynn *et al*., 2007). Whether outcomes (i) and (ii) are deleterious depends on how many crossovers remain and whether the resulting spatial distribution of crossovers usually generates at least one crossover *per* pair of homologous chromosomes (the “obligate crossover” essential for correct meiosis I segregation in almost all species (Roeder, 1997; Martini *et al*., 2006; Zanders and Alani, 2009)). Many variables could potentially reduce the deleterious impact of losing class I crossovers, but three will be discussed here: natural variation in class I and class II crossover abundance, chromosome structure, and temperature.

The proportion of class I and class II crossovers varies among different model species: nearly all *C. elegans* crossovers are lost without Msh5 (Kelly *et al*., 2000), while *ca.* 85% of *S. cerevisiae* and *A. thaliana* crossovers are class I (Borner *et al*., 2004; Higgins *et al*., 2004). It is unknown to what extent this proportion varies within lineages. If the MRCA of *A. nidulans* and *A. rugulosus* had a relatively high proportion of class II crossovers, loss of genes associated with class I crossover formation may not have been deleterious. Estimating the proportion of class I and class II crossovers in various Eurotiomycetes (particularly species in the *A. nidulans-A. conjunctus* clade) could indicate whether Eurotiomycetes as a whole have a relatively high proportion of class II crossovers (conferring a contributing predisposing “ZMM loss tolerance factor” to this group of fungi as a whole) or if species-specific modulations in the proportion of each crossover class are associated with ZMM losses.

The probability of each pair of homologous chromosomes experiencing at least one crossover can be influenced by the presence or absence of crossover interference and the associated class I crossovers. Crossover interference in *S. cerevisiae* is stronger on larger chromosomes, resulting in more widely spaced crossover events and fewer crossovers *per* kb of sequence (Kaback *et al*., 1999; Stahl *et al*., 2004). By contrast, crossovers are more densely spaced on the smaller *S. cerevisiae* chromosomes (Kaback *et al*., 1999)—the chromosomes that are predicted to be most at risk of failing to receive a crossover if all crossovers were spatially randomly distributed (Sym and Roeder, 1994) as in the Poisson distribution observed for mutants such as *A. thaliana* Msh4 knockout homozygotes (Higgins *et al*., 2004). Notably, the probability of the smallest chromosome failing to receive at least one crossover in *S. pombe* and *A. nidulans* under a Poisson distribution has been estimated to be very low (0.002-0.007% for *S. pombe*, 0.049% for *A. nidulans* (Kohli and Bahler, 1994; Sym and Roeder, 1994)), consistent with productive meiosis still occurring in these species in the absence of class I crossovers, crossover interference, Msh4, and Msh5 (Strickland, 1958; Snow, 1979; Egel-Mitani *et al*., 1982; Lynn *et al*., 2007; Malik *et al*., 2008). This probability seems to be a function of the total number of crossovers and the relative size of the smallest chromosome compared to the rest of the genome. Therefore, investigating whether taxa that have lost Msh4 and Msh5 have more crossovers or consistently different chromosome structure compared to their closest relatives retaining these genes could be illustrative (see Savelkoul, 2013 for pilot studies on chromosome size and number.)

A final variable that seems promising for future investigation is the temperature at which sexual reproduction occurs. *S. cerevisiae msh4*Δ, *msh5*Δ, *zip3*Δ, and *pch2*Δ mutants each exhibit temperature-dependent variation in phenotype severity (Borner *et al*., 2004; Chan *et al*., 2009; Joshi *et al*., 2009). For example, spore viability is less severely compromised at 33°C than at 23°C in *S. cerevisiae* SK1 and Y55 strain *msh4*Δ and *msh5*Δ mutants (Chan *et al*., 2009). This particular result would predict that species that undergo sexual reproduction primarily at warm temperatures might be predisposed to be able to better tolerate loss of Msh4 and Msh5 than species that sexually reproduce in a variety of temperature conditions or exclusively colder conditions. However, several cautions must be taken before extrapolating a universal temperature and gene loss association applicable to all species. First, strain-specific variation is evident among *S. cerevisiae msh4*Δ and *msh5*Δ phenotypes at different temperatures: the Y55 strain shows improvements in both spore viability and sporulation frequency at 33°, but the SK1 strain shows increased spore viability yet reduced sporulation frequency at 33°C compared to 23°C (Chan *et al*., 2009). Second, the more permissive temperature is not always the same for each of the mentioned *S. cerevisiae* mutants. SK1 strain *zip3*Δ mutants have improved sporulation frequency at 23°C and comparable spore viability at 23°C and 33°C (Chan *et al*., 2009), suggesting that 23°C is the overall “better” temperature for *zip3*Δ mutants. SK1 strain *pch2*Δ mutants also show less severe crossover interference defects at 30°C compared to 33°C, though crossover assurance defects in a DSB-poor background (*pch2*Δ *spo11-HA*) are more severe at 30°C (Joshi *et al*., 2009). Third, temperature effects on Msh4, Msh5, Zip3, or Pch2 knockout phenotype severity have not been extensively assessed in organisms other than *S. cerevisiae* to our knowledge to know whether temperature is a conserved influence. (An exception is a temperature-sensitive allele of the *C. elegans* Msh4 ortholog HIM-14 that is more severely affected at 23°C than 15°C (Zetka and Rose, 1995; Zalevsky *et al*., 1999).) Investigations into the effect of sexual reproduction temperature on the severity of meiosis gene losses therefore should initially proceed under the broad premise that temperature is predicted to be relevant in some respect but that the specific temperature effects may vary with lineage. With that caveat in mind, differences among thermal conditions conducive to sexual reproduction have been documented among various Eurotiomycetes. *A. nidulans* can sexually reproduce at 37°C with an optimum temperature of 32°C (Dyer, 2007 as cited in Dyer and O’Gorman, 2011; Todd *et al*., 2007; Dyer and O’Gorman, 2012), and *E. herbariorum* is known to sexually reproduce with high productivity between 27°C and 33°C (Blaser, 1975 as cited in Dyer and O’Gorman, 2012). By contrast, sexual reproduction in *A. fumigatus* and *A. flavus* under laboratory conditions requires a 30°C incubation for multiple months (Horn *et al*., 2009a; O’Gorman *et al*., 2009) and *A. terreus* requires 37°C conditions (Arabatzis and Velegraki, 2013). Interestingly, *A. nidulans* and *A. rugulosus* have a higher maximum temperature tolerance under vegetative growth conditions (48°C maximum) than their closest examined relative known to have intact Msh4 and Msh5, *A. unguis* (<40°C maximum) (Matsuzawa *et al*., 2012). Investigating whether this vegetative temperature tolerance difference is also associated with different optimal temperatures for sexual reproduction (which, based on the existence of an *Emericella unguis* teleomorph (Peterson, 2008), exists in *A. unguis* but would need to be inducible under laboratory conditions) would be the most direct test of whether temperature is indeed a relevant variable in explaining tolerance to Msh4 and Msh5 loss.

### Conclusions and Future Directions

In contrast to previous surveys of meiosis genes in Eurotiomycete fungi primarily (but not exclusively) from order Onygenales (Woo *et al*., 2006; Malik *et al*., 2008; Wang *et al*., 2009; Desjardins *et al*., 2011; Martinez *et al*., 2012), the present results identified numerous independent losses of several genes related to crossover interference and/or class I crossover formation—Msh4, Msh5, Zip3, and Pch2. Continued investigation of additional fungus species would be useful to determine whether the abundance of newly discovered meiosis gene losses is simply due to increased taxon sampling or if Eurotiomycetes are particularly prone to meiosis gene losses compared to other fungi (Savelkoul, 2013). Similarly, additional genes such as Mer3, Zip1, Zip2, and Zip4 could be investigated to see if the molecular coevolution observed among Msh4, Msh5, and Zip3 in Eurotiomycetes applies to the other “ZMM” group genes in these taxa (Lynn *et al*., 2007). The present set of Eurotiomycetes with known recent Msh4 and Msh5 losses—*A. nidulans*, *A. rugulosus*, *T. stipitatus*, and *E. herbariorum*—also represent a potentially powerful system for comparative studies of how meiosis as a whole is maintained despite losses of various meiosis genes, the extent to which convergence in other molecular traits is observed among these species, and whether these species show comparable differences in comparisons with each’s closest sister taxon retaining Msh4 and Msh5. Variables such as homothallism, chromosome size/number, the ancestral proportion of class I and class II crossovers, and temperature at sexual reproduction currently seem the most promising to investigate. Characterization of wild type meiosis properties (crossover interference or lack thereof, crossover number, synaptonemal complex formation, *etc.*) remains to be done for most Eurotiomycetes, as does functional assessment of Eurotiomycete orthologs of Msh4, Msh5, Zip3, and Pch2. However, the presence of known sexual reproduction in all four species with pseudogenized Msh4 and Msh5 (Butinar *et al*., 2005; Galagan *et al*., 2005; Todd *et al*., 2007; Peterson, 2008; Lopez-Villavicencio *et al*., 2010; Dyer and O’Gorman, 2011; Dyer and O’Gorman, 2012) as well as an increasing number of Eurotiomycetes with intact Msh4, Msh5, and Zip3—with Pch2 (*e.g., A. terreus, A. fumigatus, A. flavus* (Horn *et al*., 2009a; O’Gorman *et al*., 2009; Arabatzis and Velegraki, 2013)) or without Pch2 (*e.g., A. unguis* (Fennell and Raper, 1955))—means that comparative studies of meiosis in these Eurotiomycetes is (or may one day be) feasible. Further exploring the factors that have allowed some Eurotiomycetes to tolerate loss of otherwise well-conserved meiosis genes without losing meiosis or sexual reproduction may shed light on the variables that are most influential in constraining the evolution of sexual reproduction in Eurotiomycetes as a whole—and, potentially, other fungi.

## Materials and Methods

### Identification of Initial Eurotiomycete Query Sequences

Initial TBLASTN (Altschul *et al*., 1997) searches of the Broad Institute of MIT *Aspergillus* database (Broad, 2012; Broad, 2013), followed by identification through BLASTX comparison to the NCBI nr protein database (NCBI, 2012-2013) or BLASTP searches of the NCBI nr protein database, were performed using *S. cerevisiae* orthologs of Msh4, Msh5, Pch2 (EDV11928.1), and Zip3/Cst9 (NP_013498.3). When initial Eurotiomycete orthologs were identified (*e.g., A. clavatus*), predicted protein sequences from NCBI were typically used as queries in subsequent searches. Initial identification of Eurotiomycete Zip3 orthologs was not successful using TBLASTN *S. cerevisiae* queries and instead required PSI-BLAST (Altschul *et al*., 1997). The PSI-BLAST search, using default settings, was started with *S. cerevisiae* Cst9/Zip3 NP_013498.3, and all hits from other Saccharomycotina fungi (and no other sequences) were selected for PSSM construction. Manual inspection of the next iteration’s results identified *Trichophyton rubrum* predicted protein XP_003231222.1; BLASTX comparison to the NCBI nr protein database (NCBI, 2012-2013) found this *T. rubrum* sequence to be the reciprocal best hit of *S. cerevisiae* Cst9/Zip3.

### Bioinformatic Meiosis Gene Inventory

TBLASTN (Altschul *et al*., 1997) searches of numerous Eurotiomycete fungus genomes (*Aspergillus nidulans, A. flavus, A. niger, A. terreus, A. clavatus, A. fumigatus, Coccidioides immitis, Uncinocarpus reesii, Histoplasma capsulatum, Paracoccidioides brasiliensis, Microsporum gypseum, Microsporum canis, Trichophyton rubrum, Trichophyton tonsurans, Penicillium chrysogenum, Penicillium marneffei, Talaromyces stipitatus, Eurotium herbariorum*; see supplemental file S1 for additional genome sequence information) were performed using Eurotiomycete (Msh4, Msh5, Zip3) or *S. cerevisiae* (Pch2) predicted protein sequences with the BLOSUM62 matrix and a threshold of either E=1 (Msh4, Msh5, Zip3, Pch2) or E=0.001 (Pch2.) All initial DNA hits were retrieved with an additional 1 kbp *per* side flanking sequence for BLASTX (Altschul *et al*., 1997) comparison to the NCBI nr protein database (NCBI, 2012-2013); manual inspection of the output and/or BLink (precompiled BLASTP results) of the best scoring hit was used to identify the genomic DNA region as most similar either to the genes of interest or to non-target genes. Manual annotation of exon-intron boundaries was done in Sequencher 4.10.1 (Gene Codes Corporation) referencing BLASTX2SEQ comparisons to NCBI predictions, manual comparison of intron phases and locations, and iterative cycles of amino acid multiple sequence alignment construction in MUSCLE v3.6 (Edgar, 2004) followed by revisions. Orthologs from the above-mentioned species were subjected to later, additional phylogenetic validation of orthology (described below). The genome sequences of multiple additional Eurotiomycetes (*Aspergillus aculeatus, Aspergillus carbonarius, Aspergillus acidus, Aspergillus brasiliensis, Aspergillus glaucus, Aspergillus oryzae, Aspergillus sydowii, Aspergillus tubingensis, Aspergillus versicolor, Aspergillus wentii, Aspergillus zonatus, Coccidioides posadasii, Neosartorya fischeri, Trichophyton equinum, Trichophyton verrucosum*; supplemental file S1) were subjected to only the BLASTX and manual annotation procedures.

### Synteny Assessment

Genes flanking Msh4, Msh5, Pch2, and Zip3 orthologs in exemplar Eurotiomycetes (all four genes: *A. terreus, A. flavus, A. niger, A. clavatus, A. fumigatus, C. immitis, P. brasiliensis, U. reesii*; Msh4, Msh5, Pch2 only: *A. oryzae, H. capsulatum, M. canis, M. gypseum, N. fischeri, T. equinum, U. reesii*; Zip3 only: *P. chrysogenum* and *P. marneffei*; Figure 2) were identified by retrieving 2-5 kbp flanking sequence *per* side, subjecting those sequences to NCBI BLASTX identification as above, and manual annotation based on NCBI predictions. Predicted protein sequences of the consensus flanking genes (supplemental file S1) were used as TBLASTN queries against the *A. nidulans* genome sequence (Broad, 2013) to identify adjacent flanking gene pairs and the intergenic regions in which the four meiosis genes of interest were found in the other examined Eurotiomycetes. (Pseudogene annotation is described below in the degenerate PCR section.)

### PCR Confirmation of A. nidulans Genome Assembly

An *A. nidulans* strain A4 culture from the FGSC (McCluskey *et al*., 2010) was subcultured in *Aspergillus* minimal medium (Hill and Kafer, 2001) on 95 mm diameter plates before phenol-chloroform DNA extraction. To amplify the sequence regions corresponding to the Eurotiomycete consensus synteny for the genes flanking Msh4, Msh5, and Pch2, primary PCR with primers specific to the *A. nidulans* A4 genome sequence (Broad, 2013) was done with the following reagents *per* reaction: 10.7 µL ddH_2_O previously run through a MilliQ and autoclaved, 2.5 µL 10X Eppendorf MasterTaq buffer, 5 µL 5X Eppendorf MasterTaq buffer, 0.68 µL Fermentas/Agilent dNTP mix (40 mM total of 10mM each dNTP), 0.1 µL Eppendorf MasterTaq Taq DNA polymerase, 0.02 µL Agilent Technologies Pfu cloned polymerase, 2.5 µL each of two 10 µM Integrated DNA Technologies primers (supplemental file S1), and 1 µL of 4.2 ng/µL *A. nidulans* DNA. Primary PCR reactions were run on an Eppendorf Mastercycler gradient thermocycler for 2 minutes at 94°C; 40 cycles of denaturation (1 minute at 94°C), annealing (1 minute at 55°C ± 5°C), and extension (72°C for 2 minutes initially with 6 seconds added each cycle); and a final 10 minutes at 72°C.

To determine whether primary PCR amplification occurred, 10 µL of the PCR product solution was run on a 1% agarose gel (GeneMate LE Agarose from ISC Bioexpress) in 1X TAE buffer. If bands of the expected size were present, the remaining 15 µL of the reaction was run on a 1% to 2% agarose low-melt gel (half Fisher low melting agarose and half NuSieve agarose) in 1X TAE buffer to isolate and excise the band. PCR products within the low-melt gel slices were inserted into plasmids with the Agilent Technologies Strataclone PCR cloning kit following all manufacturer instructions except for using one-quarter-scale reactions. Isolates of the cloning kit colonies were screened for successful plasmid integration by running screening PCRs with the following reagents *per* reaction: 8.15µL sddH_2_O, 0.1 µL New England Biolabs Taq DNA polymerase, 1 µL New England Biolabs ThermoPol Buffer, 0.25 µL each of 20 µM screening PCR primers based on the Strataclone pSC-A plasmid (supplemental file S1), 0.25 µL Fermentas dNTP mix. The screening PCR program was 2 minutes at 94°C; 35 cycles of denaturation (1 minute at 94°C), annealing (1 minute at 57°C), and extension (1 minute at 72°C); and 10 minutes at 72°C. Following gel electrophoresis to confirm screening PCR amplification, cultures of positively screening colonies were grown in LB at 37°C overnight; plasmids were then isolated and purified using the Eppendorf FastPlasmid Mini kit.

Purified plasmids were sequenced using the ABI3730 with the ABI Big Dye kit and the plasmid-based sequencing primers (supplemental file S1). Sequences were analyzed in Sequencher 4.10.1 (Gene Codes Corporation) and identified by BLASTX against the NCBI nr database and BLASTN against the *A. nidulans* FGSC A4 genome sequence (Broad, 2013, also accessed 2012).

### Degenerate PCR and Pseudogene Annotation

To amplify the regions expected to contain Msh4, Msh5, Zip3, or Pch2 from taxa without available whole genome sequences, degenerate PCR primers were designed based on conserved amino acid sequences within the corresponding flanking gene pairs for each meiosis gene based on the Eurotiomycete consensus synteny for these regions. Previously extracted DNA from *Emericella rugulosa (Aspergillus rugulosus)* NRRL 206, *Aspergillus unguis* NRRL 216, *Aspergillus versicolor* NRRL 238, *Aspergillus crustosus* NRRL 4988, *Aspergillus ustus* NRRL 4991, *Aspergillus funiculosus* NRRL 4744, *Aspergillus conjunctus* NRRL 5080, *Aspergillus penicillioides* NRRL 4548, and *Aspergillus clavatoflavus* NRRL 5113 was graciously provided by Dr. Stephen Peterson of the USDA; DNA stocks were diluted to 3 ng/µL for PCR. Primary degenerate PCR was initially done as described above for the *A. nidulans* confirmation; in some cases, nested and/or hemidegenerate PCR was used (primers in supplemental file S1.) Later reactions used different reagents (*per* reaction: 2.75 µL Promega GoTaq MgCl_2_, 7.75 µL green 5X Promega GoTaq Flexi Buffer, 0.19 µL Promega GoTaq polymerase, 0.93 µL of Fermentas 40 mM dNTP mix [10 mM *per* dNTP], 0.03 µL Agilent Technologies Pfu cloned polymerase, 23.75 µL ddH_2_O previously run through a MilliQ and autoclaved, 0.75 µL of each 50 µM primer from Integrated DNA Technologies (supplemental file S1)) and a non-gradient PCR program (2 minutes at 94°C; 10 cycles of 30 seconds at 94°C, 1 minute at 50°C, 5 minutes at 72°C; 30 cycles of 30 seconds at 94°C, 1 minute at 50°C, 5 minutes plus an additional 10 seconds each cycle at 74°C; 10 minutes at 74°C). Gel electrophoresis, cloning, and sequencing was performed as above for *A. nidulans* with two exceptions: first, some samples used the Zymo Research Zyppy Plasmid Miniprep Kit for plasmid purification prior to sequencing; second, if initial sequencing reads covered only the 5’ and 3’ ends of the region, additional sequence-specific primers (supplemental file S1) were used to extend the sequence.

BLASTX identification and manual annotation of sequenced PCR products was done as described for the bioinformatic meiosis gene inventory section with a few exceptions. Some regions that were amplified with only 1X local coverage had places where a single “N” placeholder base was needed to maintain an otherwise highly conserved reading frame; since these were consistent with a single indel sequencing error due to low coverage and the surrounding sequence was highly conserved across Eurotiomycetes, we did not interpret these cases (single “N” placeholder in 1X local coverage) as pseudogenes. To be included in the phylogenetic analysis, the putative pseudogenes (sequences with multiple putative frameshift mutations) required introduction of “N” placeholder bases to be able to be included in the amino acid multiple sequence alignment. All “N” placeholders were added to the first or second codon position whenever possible for the predicted residue to be an ambiguous “X” that would not inappropriately speculate on the former residue encoded at that site.

### Phylogenetic Analysis

Amino acid multiple sequence alignments of Msh4, Msh5, Zip3, and Pch2 were each constructed in MUSCLE v3.6 (Edgar, 2004) under default settings. Regions that were poorly aligned were manually excised using Se-Al v2.0a11 (Rambaut, 2002) to use only unambiguously aligned sites; MEGA5.05 (Tamura *et al*., 2011) was then used to determine the optimal sequence substitution model for each alignment (default settings, all sites in trimmed alignment considered). Each maximum likelihood phylogeny was constructed in PhyML v3.0 (Guindon *et al*., 2010) on the ATGC: Montpelier Bioinformatics platform (Lefort, 2013, also accessed 2012) using a JTT+G5 model (Yang, 1994; Jones *et al*., 1992), the “best of NNI and SPR” search method (Guindon and Gascuel, 2003; Guindon *et al*., 2010), and 1000 bootstrap replicates (Felsenstein, 1985). Amino acid multiple sequence alignments without manual trimming will be in supplemental files S2-S5 (available on request).

### Topology Tests

The relationship of *A. penicillioides* and *A. clavatoflavus* relative to the major Pch2-retaining *Aspergillus* clade (*A. fumigatus*-*A. terreus*) and Pch2-lacking *Aspergillus* clade (*A. nidulans-A. conjunctus*) was assessed in TREE-PUZZLE v5.2 (Schmidt *et al*., 2002). Twelve topologies varying the placement of *A. penicillioides* and *A. clavatoflavus* relative to the two known clades (Figure 9) were compared utilizing six amino acid alignment inputs: Msh4 alone, Msh4 concatenated with available flanking gene sequence, Msh5, Msh5 concatenated with available flanking gene sequence, concatenated Msh4 and Msh5, and concatenated Msh4 and Msh5 with available flanking gene sequences. (The Msh4 and Msh5 alignments were taken from the phylogenetic analyses, while the flanking gene sequences include only regions amplified from the degenerate PCR primers, *i.e.,* full-length flanking gene sequences were not utilized for species with sequenced genomes.) TREE-PUZZLE utilized four tests (1sKH and 2sKH: one-sided or two-sided Kishino-Hasegawa tests based on pairwise SH tests (Kishino and Hasegawa, 1989; Goldman *et al*., 2000), SH: Shimodaira-Hasegawa test (Shimodaira and Hasegawa, 1999), ELW: expected likelihood weight (Strimmer and Rambaut, 2002)); all four tests were considered in evaluating whether a particular topology could be rejected as significantly worse than the optimal topology based on comparisons of log likelihood (log L) values of each topology given the input alignment.

### Codeml Analysis of K_a_/K_s_ in Aspergillus Msh4, Msh5, and Zip3 Relative to Pch2 Loss

The PAML program Codeml (Yang, 2007) was used under default settings to test whether models assuming a different underlying K_a_/K_s_ ratio for any or all Pch2-lacking *Aspergillus* species (variants of the model=2 setting) were significantly better than null models using a likelihood ratio test. To ensure that reading frames were maintained in the DNA alignments, amino acid multiple sequence alignment information was maintained, TranslatorX (Abascal *et al*., 2010) was used to generate a DNA alignment based on the MUSCLE (Edgar, 2004) amino acid multiple sequence alignments; sections that were ambiguously aligned and removed from the amino acid alignments beforeML phylogenetic analysis were also removed from the DNA alignment. All topologies had the *A. nidulans-A. conjunctus* clade sister to the *A. fumigatus-A. terreus* clade (topology 1 in Figure 9, see Figure 1 for relationships within noted clades). The outgroup was either *U. reesii* with *A. clavatoflavus* and *A. penicillioides* included (Table 2A-2B, Table 3A-3B), *A. penicillioides* (Table 2C-2D, Table 3C-3D), or *P. chrysogenum* (Table 4). Likelihood ratio tests were used to assess whether compared models were significantly different from each other.

### Codeml Analysis of Zip3 K_a_/K_s_ in Eurotiomycetes with Pseudogenized Msh4 and Msh5

A core set of species with intact Zip3 (*A. versicolor, A. conjunctus, A. crustosus, A. unguis, A. clavatus, A. fumigatus, A. flavus, A. terreus, A. niger, P. chrysogenum;* outgroup: *Uncinocarpus reesii*) along with four different combinations of additional species (*E. herbariorum*, *P. marneffei*, *T. stipitatus*, *P. marneffei* and *T. stipitatus*) was used to determine whether Zip3 K_a_/K_s_ is significantly different in species with pseudogenized Msh4 or Msh5. Codeml (Yang, 2007) analyses and DNA alignment construction was performed as above. Although *E. herbariorum*’s phylogenetic relationship to the major *Aspergillus* clades was an unresolved polytomy in the work of Peterson (2008), that work showed strong support for *E. herbariorum* and *A. penicillioides* forming a clade (Peterson, 2008); therefore, *E. herbariorum* here was put in the *A. penicillioides* position of topology 1 in Figure 9. *P. chrysogenum*, *P. marneffei*, and *T. stipitatus* were added to topology 1 according to previous findings (van den Berg *et al*., 2008).

### Identification of Putative MAT Gene Orthologs in Exemplar Eurotiomycetes

Exemplar *A. fumigatus* MAT1-1-1 and MAT1-2-1 protein sequences were obtained by NCBI keyword searches for “*Aspergillus”* and “MAT1-1-1” or “MAT1-2-1” (accession numbers listed in Figure 12). Genome sequence TBLASTN searches followed by NCBI nr protein database BLASTX identification was performed as described in the “Bioinformatic Meiosis Gene Inventory” subsection for the species shown in Figure 12.

## Acknowledgements

This work was supported by the Carver Trust, the University of Iowa Department of Biology, and the University of Iowa Graduate College. We thank the Fungal Genetics Stock Center (FGSC) for *A. nidulans* cultures and Stephen Peterson (USDA) for providing various *Aspergillus* DNA samples. Nathan Benassi (University of Iowa Honors undergraduate) performed *A. nidulans* genome validation PCR, and Sneha Patil (University of Iowa Secondary Student Training Program) contributed to some of the Zip3 degenerate PCR work. We thank the Broad Institute of MIT, the U.S. Department of Energy Joint Genome Institute, and NCBI for providing and allowing public use of the examined Eurotiomycete genome sequences. We gratefully acknowledge the original submitters of the many predicted protein sequences deposited in the NCBI nr protein database; the accession numbers for specific sequences are provided in supplemental file S1 and, for MAT genes, in Figure 12. Thank you to James Gloer, Robert Malone, Maurine Neiman, and David Soll for providing helpful feedback on an earlier version of this work in a thesis dissertation context.

## Supplementary Information

Supplemental File S1 (.xlsx): (A) Genome data, (B) accession numbers, (C) primers.

*\*Supplemental File S2: Untrimmed Msh4 alignment*

*\*Supplemental File S3: Untrimmed Msh5 alignment*

*\*Supplemental File S4: Untrimmed Pch2 alignment*

*\*Supplemental File S5: Untrimmed Zip3 alignment*

*\*Note: these supplemental files are available on request*.

